# The genome of cowpea (*Vigna unguiculata* [L.] Walp.)

**DOI:** 10.1101/518969

**Authors:** Stefano Lonardi, María Muñoz-Amatriaín, Qihua Liang, Shengqiang Shu, Steve I. Wanamaker, Sassoum Lo, Jaakko Tanskanen, Alan H. Schulman, Tingting Zhu, Ming-Cheng Luo, Hind Alhakami, Rachid Ounit, Abid Md. Hasan, Jerome Verdier, Philip A. Roberts, Jansen R.P. Santos, Arsenio Ndeve, Jaroslav Doležel, Jan Vrána, Samuel A. Hokin, Andrew D. Farmer, Steven B. Cannon, Timothy J. Close

## Abstract

Cowpea (*Vigna unguiculata* [L.] Walp.) is a major crop for worldwide food and nutritional security, especially in sub-Saharan Africa, that is resilient to hot and drought-prone environments. A high-quality assembly of the single-haplotype inbred genome of cowpea IT97K-499-35 was developed by exploiting the synergies between single molecule real-time sequencing, optical and genetic mapping, and a novel assembly reconciliation algorithm. A total of 519 Mb is included in the assembled sequences. Nearly half of the assembled sequence is composed of repetitive elements, which are enriched within recombination-poor pericentromeric regions. A comparative analysis of these elements suggests that genome size differences between *Vigna* species are mainly attributable to changes in the amount of *Gypsy* retrotransposons. Conversely, genes are more abundant in more distal, high-recombination regions of the chromosomes; there appears to be more duplication of genes within the NBS-LRR and the SAUR-like auxin superfamilies compared to other warm-season legumes that have been sequenced. A surprising outcome of this study is the identification of a chromosomal inversion of 4.2 Mb among landraces and cultivars, which includes a gene that has been associated in other plants with interactions with the parasitic weed *Striga gesnerioides*. The genome sequence also facilitated the identification of a putative syntelog for multiple organ gigantism in legumes. A new numbering system has been adopted for cowpea chromosomes based on synteny with common bean (*Phaseolus vulgaris*).

## INTRODUCTION

Cowpea (*Vigna unguiculata* [L.] Walp.) is one of the most important food and nutritional security crops, providing the main source of protein to millions of people in developing countries. In sub-Saharan Africa, smallholder farmers are the major producers and consumers of cowpea, which is grown for its grains, tender leaves and pods as food for human consumption, with the crop residues being used for fodder or added back to the soil to improve fertility^1^. Cowpea was domesticated in Africa, from where it spread into all continents and now is commonly grown in many parts of Asia, Europe, the United States, and Central and South America. One of the strengths of cowpea is its high resilience to harsh conditions, including hot and dry environments, and poor soils. Still, as sub-Saharan Africa and other cowpea production regions encounter climate variability^2, 3^, breeding for more climate-resilient varieties remains a priority.

Cowpea is a diploid member of the Fabaceae family with a chromosome number 2n = 22 and a previously estimated genome size of 613 Mb^4^. Its genome shares a high degree of collinearity with other warm season legumes (Phaseoleae tribe), including common bean (*Phaseolus vulgaris* L.)^5, 6^. A highly fragmented draft assembly and BAC sequence assemblies of IT97K-499-35 were previously generated^5^. Although these resources enabled progress on cowpea genetics and genomics^7–11^, they lacked the contiguity and completeness required for accurate genome annotation, detailed investigation of candidate genes or thorough genome comparisons. Here, we re-estimated the genome size of *Vigna unguiculata* by k-mer analysis and flow cytometry, and produced a high-quality genome assembly using single-molecule real-time sequencing and optical and genetic mapping. This cowpea reference sequence has been used for the analysis of repetitive elements, gene families, and genetic variation, and for comparative analysis with three close relative legumes including common bean, which stimulated a change of chromosome numbering to facilitate comparative studies. The publicly available genome sequence of cowpea lays the foundation for all forms of basic and applied research, enabling progress towards the genetic improvement of this key crop plant for food and nutritional security.

## RESULTS AND DISCUSSION

### Estimation of *Vigna unguiculata* genome size using flow cytometry and k-mer distribution

To estimate the genome size of the sequenced reference accession IT97K-499-35 (see below), nuclear DNA content was estimated using flow cytometry^12^ and k-mer analysis. In brief, the cytometry results indicated that the 2C nuclear DNA amount of *Vigna unguiculata* IT97K-499-35 is 1.310 ± 0.026 pg DNA (mean ± SD), which corresponds to 1C genome size of 640.6 Mbp. This is higher than the previous estimate of 613 Mbp by Arumuganathan and Earle^4^, but 841 Mbp smaller than the estimate of Parida et al.^13^. Estimate differences could be due to method/protocol dissimilarities, or to other factors including instrument variation between laboratories and actual differences between the accessions analyzed (see Methods). As is commonly done in large-scale sequencing projects, a k-mer distribution analysis was also carried out. The k-mer distribution analysis (k=27) using ~168M 149-bp paired-end Illumina reads yielded an estimated genome size of 560.3 Mbp (see Methods and Supplementary Figure 1).

**Figure 1.**
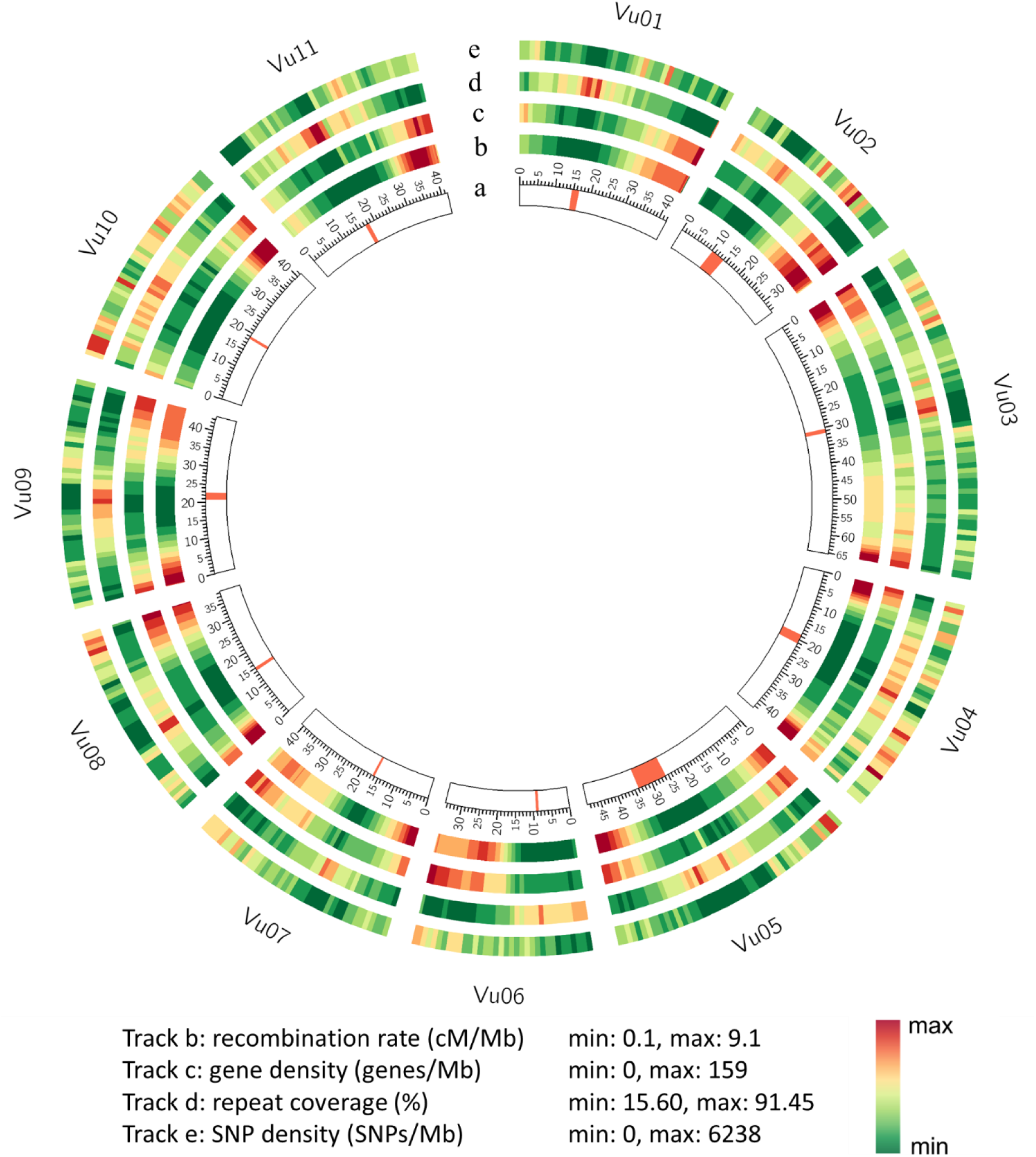
Landscape of the cowpea genome. (a) Cowpea chromosomes in Mb, with red lines representing centromeric regions based on a 455-bp tandem repeat alignment^26^; (b) Recombination rate at each 1Mb; (c) Gene density in 1Mb windows; (d) Repeat coverage in 1Mb windows; (e) SNP density in 1Mb windows.

### Sequencing and assembly using a novel “stitching method”

The elite breeding line IT97K-499-35, developed at the International Institute of Tropical Agriculture (IITA, Nigeria), was used previously for the development of genome resources^5,14^. Here, a fully homozygous (single haplotype; See Methods) stock was sequenced using PacBio (Pacific Biosciences of California, Inc., Menlo Park, CA, USA) single-molecule real-time (SMRT) sequencing. In total, 56.8 Gb of sequence data were generated (~91.7x genome equivalent), with a read N50 of 14,595 bp. Pre-and post-filter read length and quality distribution are reported in Supplementary Figures 2-4. Also, two Bionano Genomics (San Diego, California, USA) optical maps^15^ were generated using nicking enzymes *BspQ*I and *BssS*I (Supplementary Tables 1 and 2). The size of the *BsqQ*I optical map is 622.21 Mb, while the size of the *BssS*I optical map is 577.76 Mb.

With the PacBio data, eight draft assemblies were generated, six of which were produced with CANU^16, 17^ using multiple parameter settings at the error correction stage, one with Falcon^18^, and one with ABruijn^19^. As Supplementary Table 3 shows, CANU, Falcon, and ABruijn produced assemblies with significantly different assembly statistics, which made it difficult to designate one as “best”. These tools are fundamentally different at the algorithmic level (e.g., CANU and Falcon are based on the overlap-layout-consensus paradigm, while ABruijn uses the de Bruijn graph), and their designers have made different choices in the tradeoff between maximizing assembly contiguity (i.e., N50) versus minimizing the probability of mis-assemblies (i.e., mis-joins). Here, we employed an alternative assembly methodology: instead of having to choose one of the assemblies, one or more optical maps were leveraged to merge multiple assemblies (see Pan et al.^20^ and Methods for details) in what we called “stitching.” This method was applied to the eight assemblies in Supplementary Table 3, after removing contaminated contigs and breaking chimeric contigs identified using the optical maps. The number of chimeric contig ranged from 16 to 40 depending on the assembly. Each of the eight assemblies contributed a fraction of its contigs to the final assembly: 13% of the “minimal tiling path” (MTP) contigs were from the FALCON assembly, 8% from the ABruijn assembly and the rest (79%) from the six CANU assemblies each ranging from 4% to 20%. Table 1 reports the assembly statistics of the stitched and polished (PacBio Quiver pipeline) assembly. PacBio Quiver enables consensus accuracies on genome assemblies approaching or exceeding Q60 (one error per million bases) when the sequencing depth is above 60X^92^. Note that all of the assembly statistics significantly improved compared to the eight individual assemblies (Supplementary Table 3). For instance, the N50 for the stitched assembly (10.9 Mb) was almost double the best N50 for any of the eight individual assemblies. Similarly, the longest contig for the stitched assembly increased by 4 Mb over the longest contig of any single assembly.

**Table 1.** Assembly statistics for stitched contigs, scaffolds, and pseudomolecules.

|  | Stitched contigs | Scaffolds | Pseudomolecules |
| --- | --- | --- | --- |
| N50 (bp) | 10,911,736 | 16,417,655 | 41,684,185 |
| L50 | 16 | 12 | 6 |
| NG50 (bp) | 9,203,620 | 15,388,583 | 41,327,797 |
| LG50 | 21 | 15 | 7 |
| total (bp) | 518,799,885 | 519,432,264 | 519,435,864 |
| contigs/scaffolds | 765 | 722 | 686 |
| contigs/scaffolds ≥ 100kbp | 177 | 135 | 103 |
| contigs/scaffolds ≥ 1Mbp | 61 | 38 | 13 |
| contigs/scaffolds ≥ 10Mp | 18 | 21 | 11 |
| longest contig/scaffold (bp) | 22,343,392 | 30,539,429 | 65,292,630 |
| % N | 0.0% | 0.523% | 0.524% |
| mapped SNPs | 49,888 | 49,888 | 49,888 |
| GC | 33.0% | 32.994% | 32.994% |

Scaffolds were obtained by mapping the stitched and polished assembly to both optical maps using the Kansas State University pipeline^21^. Briefly, a total of 519.4 Mb of sequence scaffold were generated with an N50 of 16.4 Mb (Table 1). Finally, a total of ten genetic maps containing 44,003 unique Illumina iSelect SNPs^5^ were used to anchor and orient sequence scaffolds into eleven pseudo-molecules via ALLMAPS^22^. Details of the ten genetic maps can be found in Supplementary Table 4. ALLMAPS was able to anchor 47 of the 74 scaffolds for a total of 473.4 Mb (91.1% of the assembled sequences), 30 of which were also oriented, resulting in 449 Mb of anchored and oriented sequence (Table 1). Only 46 Mb (8.9% of the total assembly) were unplaced. The average GC content of the assembly was 32.99%, similar to other sequenced legumes^23–25^. The quality of the chromosome-level assembly was evaluated using a variety of metrics. Several sequence datasets that were independently generated were mapped onto the assembly using BWA-mem with default settings, namely (1) about 168M 149-bp paired-end Illumina reads (98.92% mapped of which 86.7% were properly paired and 75.53% had MAPQ of at least 30), (2) about 129 thousand contigs (500 bp or longer) of the WGS assembly generated previously^5^ (99.69% mapped of which 98.69% had MAPQ>30), (3) about 178 thousand BAC sequence assemblies generated previously^5^ (99.95% mapped of which 68.39% had MAPQ>30), and (4) about 157 thousand transcripts generated for study^27^ (99.95% mapped of which 94.74% had MAPQ>30). All of these metrics indicate agreement with the pseudo-molecules at the nucleotide level.

The original PacBio reads were also mapped onto the assembly using BLASR using default settings: 5.29M long reads mapped for a total of about 46×10^9^ bp. Based on the fact that 88.68% of the bases of the long reads were present in the assembly, the cowpea genome size can be re-estimated at 585.8 Mbp, which is intermediate between the new k-mer-and cytometry-based estimates (see above).

### New chromosome numbering for cowpea

Several members of the Phaseoleae tribe are diploid with 2n = 22, but the numbering of chromosomes has been designated independently by each research group. Among these species, the *P. vulgaris* genome sequence was the first to be published^23^, thus establishing a precedent and rational basis for a more uniform chromosome numbering system within the Phaseoleae. Extensive synteny has been previously observed between cowpea and common bean^5^. By virtue of the more complete reference genome sequence, cowpea can now serve as a convenient nodal species for this cross-genus comparison, including a new chromosome numbering system based on synteny with common bean.

As summarized in Supplementary Figure 6 and Supplementary Table 5, six cowpea chromosomes are largely syntenic with six common bean chromosomes in one-to-one relationships, making the numbering conversion straightforward in those cases. Each of the remaining five cowpea chromosomes is related to parts of two *P. vulgaris* chromosomes (one-to-two relationships). For each of those cases, we adopted the number of the common bean chromosome sharing the largest syntenic region with cowpea. There was one exception to this rule: two cowpea chromosomes (previous linkage groups/chromosomes #1 and #5) both shared their largest block of synteny with *P. vulgaris* chromosome (Pv08). However, there was only one optimum solution to the chromosome numbering of cowpea, assigning the number 8 to previous cowpea linkage group/chromosome #5 and assigning the number 5 to previous linkage group/chromosome #1 (Supplementary Table 5).

In addition, comparisons between cowpea genetic maps and chromosomal maps developed by fluorescence in situ hybridization (FISH) using cowpea BACs as probes^26^ revealed that the prior orientations of three linkage groups (now referred to as Vu06, Vu10, and Vu11) were inverted relative to their actual chromosome orientation. Hence, cowpea pseudomolecules and all genetic maps were inverted for chromosomes Vu06, Vu10, and Vu11 to meet the convention of short arm on top and long arm on the bottom, corresponding to ascending cM values from the distal (telomeric) end of the short arm through the centromere and on to the distal end of the long arm. The new numbering system is shown in Supplementary Table 5 and used throughout the present manuscript. The Windows software HarvEST:Cowpea (harvest.ucr.edu), which includes a synteny display function, also has adopted this new numbering system.

### Gene annotation and repetitive DNA

The assembled genome was annotated using *de novo* gene prediction and transcript evidence based on cowpea ESTs^63^ and RNA-seq data from leaf, stem, root, flower and seed tissue^11,27^ and protein sequences of Arabidopsis, common bean, soybean, Medicago, poplar, rice and grape (see Methods). In total, 29,773 protein-coding loci were annotated, along with 12,514 alternatively spliced transcripts. Most (95.9%) of the 1,440 expected plant genes in BUSCO v3^28^ were identified in the cowpea gene set, indicating completeness of genome assembly and annotation. The average gene length was 3,881 bp, the average exon length was 313 bp, and there were 6.29 exons per gene on average. The GC content in coding exons was higher than in introns + UTRs (40.82% vs. 24.27%, respectively). Intergenic regions had an average GC content of 31.84%.

Based on the results of an automated repeat annotation pipeline (Supplementary Table 6), an estimated 49.5% of the cowpea genome is composed of the following repetitive elements: 39.2% transposable elements (TEs), 4% simple sequence repeats (SSRs), and 5.7% unidentified low-complexity sequences. The retrotransposons, or Class I TEs, comprise 84.6% of the transposable elements by sequence coverage and 82.3% by number. Of the long terminal repeat (LTR) retrotransposons, elements of the *Gypsy* superfamily^29^ (code RLG) are 1.5 times more abundant than *Copia* (code RLC) elements, but non-autonomous TRIM elements appear to be very rare, with only 57 found. The LINEs (RIX) and SINEs (RSX), comprising the non-LTR retrotransposons, together amount to only 0.4% of the genome. The DNA, or Class II, transposons compose 6.1% of the genome, with the CACTA (DTC; 5.7% of the transposable element sequences), hAT (DTA; 3.5%), and MuDR (DTM; 2.4%) being the major groups of classical “cut-and-paste” transposons. The rolling-circle *Helitron* (DHH) superfamily is relatively abundant at 1.3% of the genome and 7013 individual elements. Only 6.4% of the TE sequences were unclassified.

Centromeric regions were defined based on a 455-bp tandem repeat that was previously identified by FISH as abundant in cowpea centromeres^26^. Regions containing this sequence span over 20.18 Mb (3.9% of the assembled genome; Supplementary Table 7). Cowpea centromeric and pericentromeric regions are highly repetitive in sequence composition and exhibit low gene density and low recombination rates, while both gene density and recombination rate increase as the physical position becomes more distal from the centromeres (Figure 1; Supplementary Figure 7; Supplementary Table 8). Contrasting examples include Vu04, where the recombination rate near the telomeres of both arms of this metacentric chromosome are roughly ten times the rate across the pericentromeric region, versus Vu02 and Vu06, where the entire short arm in each of these acrocentric chromosomes has a low recombination rate (Supplementary Figure 7). These patterns have been observed in other plant genomes including legumes^23, 30^ and have important implications for genetic studies and plant breeding. For example, a major gene for a trait that lies within a low recombination region can be expected to have a high linkage drag when introgressed into a new background. As a consequence, knowledge of the recombination rate can be integrated into decisions on marker density and provide weight factors in genomic selection models to favor rare recombination events within low recombination regions.

### Cowpea genetic diversity

#### Single-nucleotide and insertion/deletion variation

Whole-genome shotgun (WGS) data from an additional 36 diverse accessions relevant to Africa, China and the USA were previously used to identify single nucleotide polymorphisms^5^. Almost all (99.83%) of the 957,710 discovered SNPs (hereinafter referred as the “1M list”) were positioned in the reference sequence, including 49,697 SNPs that can be assayed using the Illumina iSelect Consortium Array^5^ (Supplementary Table 9). About 35% of the SNPs in the 1M list were associated with genes (336,285 SNPs), while that percentage increased to 62% in the iSelect array (31,708 SNPs; Supplementary Tables 9 and 10). This indicates that the intended bias towards genes in the iSelect array design^5^ was successful. The number of cowpea gene models containing SNPs was 23,266 (78%), or 27,021 (91% of annotated genes) when considering genes at a distance of <10 kb from a SNP (Supplementary Table 10). In general, SNP density was lowest near the centromeric regions (Figure 1), including some blocks lacking SNPs entirely (Supplementary Figure 8). The SNP information provided here enables formula-based selection of SNPs, including distance to gene and recombination rate. When these metrics are combined with minor allele frequency and nearness to a trait determinant, one can choose an optimal set of SNPs for a given constraint, for example cost minimization, on the number of markers.

The same WGS data described above were analyzed using BreakDancer v.1.4.5^31^ to identify structural variants relative to the reference genome, IT97K-499-35. A total of 17,401 putative insertions and 117,403 putative deletions were identified (Supplementary Table 11). The much smaller number of insertions than deletions may reflect limitations in the ability of the software to identifying insertions when sequence reads are mapped to a reference genome. To check the fidelity of these putative variants, deletions that were apparent from the iSelect SNP data available from those 36 accessions were used. If a deletion really exists in a particular accession, then any SNP assayed within the deleted region must yield a “NoCall” from the iSelect assay. Among the 5,095 putative deletions that spanned SNPs represented in the iSelect array, only 1,558 (30.6%) were validated by this method. This indicates that the presently available data from one reference-quality genome sequence and WGS short reads from 36 accessions are insufficient to create a comprehensive or reliable catalog of structural variants; additional high-quality *de novo* assemblies are needed for reliable identification of structural variants in cowpea and will be the subject of work to be described elsewhere.

#### Identification of a 4.2 Mb chromosomal inversion on Vu03

As explained above, a total of ten genetic maps were used to anchor and orient scaffolds into pseudomolecules. Plots of genetic against physical positions for SNPs on seven of those genetic maps showed a relatively large region in an inverted orientation (Figure 2A; Supplementary Figure 9). The other three genetic maps showed no recombination in this same region, suggesting that the two parents in the cross had opposite orientations. A closer look at the genotype data from all of the parental lines showed that one of the parents from each of those three populations had the same haplotype as IT97K-499-35, and hence they presumably the same orientation (Supplementary Table 12). To validate this inversion and define inversion breakpoints, available WGS data from some of these accessions^5^ were used. In both breakpoint regions, contigs from accessions that presumably had the same orientation as the reference (type A) showed good alignments, while those from accessions with the opposite orientation (type B) aligned only until the breakpoints (Supplementary Table 13). An additional *de novo* assembly of a ‘type B’ accession (unpublished) enabled a sequence comparison with the reference genome for the entire genomic region containing the inversion (Figure 2B). This provided a confirmation of the chromosomal inversion and the position of the two breakpoints in the reference sequence: 36,118,991 bp (breakpoint 1) and 40,333,678 bp (breakpoint 2) for a 4.21 Mb inversion containing 242 genes (Supplementary Table 14). PCR amplifications of both breakpoint regions further validated this inversion (see Methods and Supplementary Figure 10).

**Figure 2.**
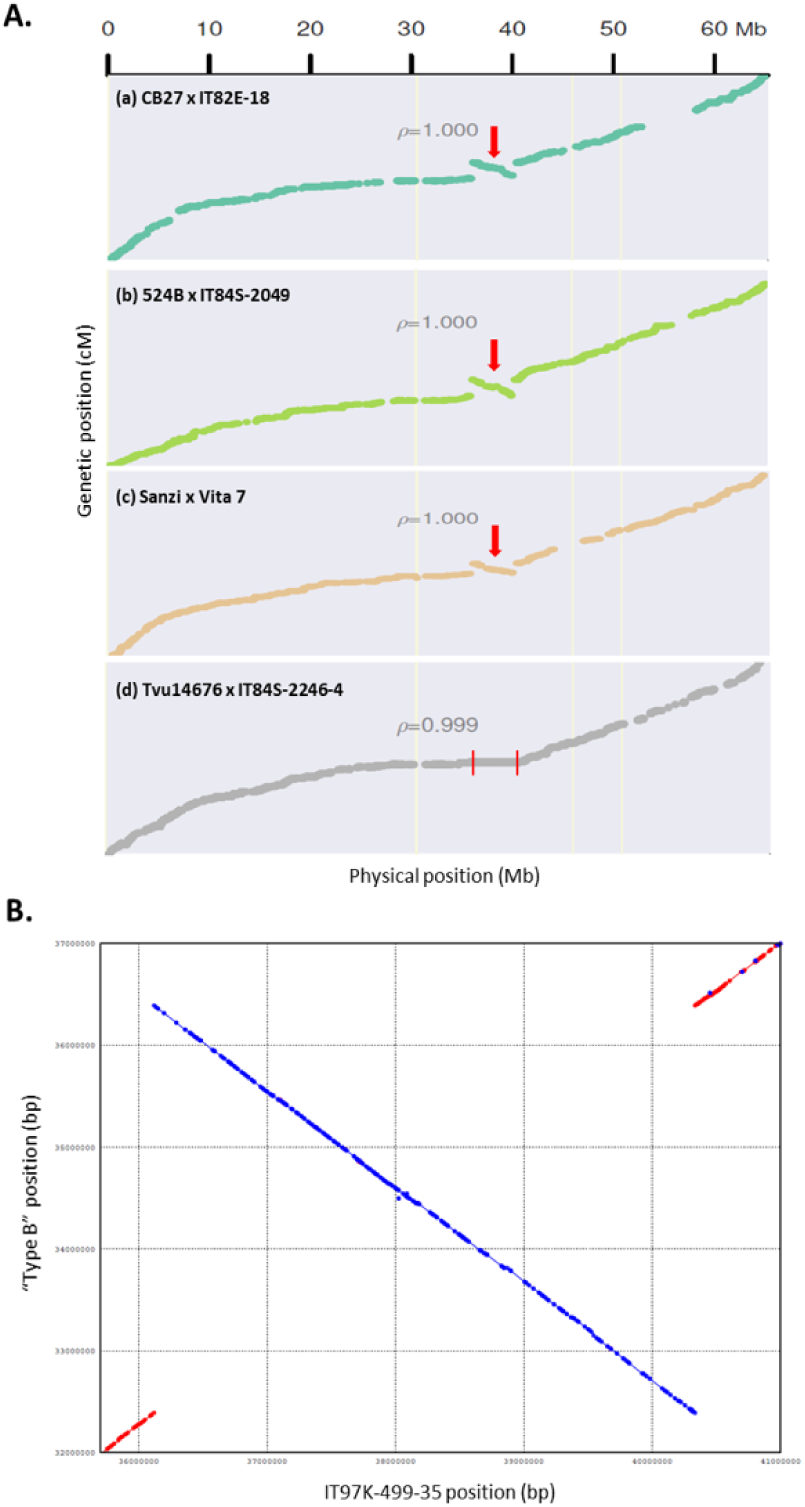
Large chromosomal inversion detected on Vu03. (A) The relationships between genetic and physical positions are shown for SNPs on four genetic maps (a to d). Maps (a) to (c) show a 4.2 Mb region in an inverted orientation (red arrow) while map (d) shows no recombination in that same region (area contained within red lines). (B) Sequence comparison between IT97K-499-35 (reference genome) and a “type B” accession for the region including the Vu03 chromosomal inversion. Red color indicates same orientation between both sequences, while in blue are shown those sequences having opposite orientations between accessions.

A set of 368 diverse cowpea accessions, including 243 landrace and 97 breeding accessions, for which iSelect data existed was used to estimate the frequency of the inversion among germplasm accessions. A total of 33 accessions (9%) had the same SNP haplotype as the reference genome across the entire region, which we presume to indicate the same orientation. Among those 33 accessions, only three were landraces (1.2% of the landraces in the set), while the other 30 were breeding materials, including the reference genome. This suggests that the orientation of this region in the reference genome is the less common orientation. Also, a complete lack of recombination across this region is reflected in the genetic map derived from a cultivated × wild cross^9^ (IT99K-573-1-1 × TVNu-1158; Supplementary Figure 9), which indicates that the wild parent has the opposite orientation of the cultivated accession. Since this cultivated parent has the same haplotype as the reference genome, and thus presumably also the same orientation, the lack of recombination across this region suggests that the opposite-to-reference orientation is the ancestral (wild) type while the reference orientation carries an inversion. A comparison between cowpea and adzuki bean for the inversion region using a gene-family assignment strategy (Supplementary Figure 11) showed that IT97K-499-35 and adzuki bean genome assemblies have opposite orientations, consistent with the conjecture that the cowpea reference genome is inverted in this region with respect to an ancestral state that has been retained in the wild cowpea accession as well as in this representative congeneric species.

A direct effect of inversions is that they suppress recombination in heterozygotes, causing ancestral and inverted types to evolve independently. Selection can act to maintain an inversion when it carries one or more advantageous alleles or when an inversion breakpoint causes gene disruption or expression changes that are adaptive^32, 33^. Two of the three landraces carrying the inversion (B-301 and B-171) originated from Botswana while the third (TVu-53) is a Nigerian landrace. Interestingly, B-301 was the key donor of resistance to several races of *Striga gesnerioides*, a serious parasitic weed of cowpea, and is in the pedigree of many breeding lines that carry the inversion, most of which are also *Striga* resistant (including the reference genome IT97K-499-35). To explore whether the inversion is associated with *Striga* resistance, the map positions of previously identified QTLs for this trait^34–36^ were compared to the position of the inversion. QTLs for resistance to *Striga* Races 1 and 3 were located on a different chromosome/linkage group than the inversion on Vu03, ruling out the inversion for those resistances. However, it is also possible that inversion breakpoints or orientation could disrupt or alter the expression of a gene involved in *Striga* interactions, such as strigolactone production, for example. Such changes could affect *Striga* germination, as in sorghum^37^. The gene annotations are not certain at the insertion break points. However, it was noted that the sorghum gene *Sobic.005G213600* regulating *Striga* resistance via a presence/absence variation^37^ encodes a sulfotransferase that is homologous to the cowpea gene *Vigun03g220400*, which is located inside the inverted region on Vu03 (Supplementary Table 14) and is highly expressed in root tissue (https://legumeinfo.org/feature/Vigna/unguiculata/gene/vigun.IT97K-499-35.gnm1.ann1.Vigun03g220400#pane=geneexpressionprofile). Therefore, it seems possible that the relocation of *Vigun03g220400* as a consequence of the inversion may have altered its position relative to regulatory elements, thus affecting its expression and *Striga* interactions in a manner that has not yet been discovered. This hypothesis merits further testing. In addition to *Striga* considerations, a QTL for pod number^38^ (*Qpn.zaas-3*) is located inside the inverted region.

Even though additional studies will be required determine whether there is an adaptive advantage for the Vu03 inversion, awareness of it is important for trait introgression and breeding, as this region represents nearly 1% of the cowpea genome and can be moderately active recombinationally during meiosis only when both chromatids carry the same orientation.

### Synteny with other warm season legumes

Synteny analyses were performed between cowpea and its close relatives adzuki bean (*Vigna angularis*), mung bean (*Vigna radiata*) and common bean (*Phaseolus vulgaris*). Extensive synteny was observed between cowpea and the other three diploid warm-season legumes although, as expected, a higher conservation was observed with the two *Vigna* species (Figure 3A-C) than with common bean. Six cowpea chromosomes (Vu04, Vu06, Vu07, Vu09, Vu10 and Vu11) largely have synteny with single chromosomes in all three other species. Cowpea chromosomes Vu2, Vu03 and Vu08 also have one-to-one relationships with the other two *Vigna* species but one-to-two relationships with *P. vulgaris*, suggesting that these chromosome rearrangements are characteristic of the divergence of *Vigna* from *Phaseolus*. The remaining cowpea chromosomes Vu01 and Vu05 have variable synteny relationships, each with two chromosomes in each of the other three species, suggesting these chromosome rearrangements are more characteristics of speciation within the *Vigna* genus. It should be noted also that most chromosomes that have a one-to-two relationship across these species or genera are consistent with translocations involving the centromeric regions (Figure 3A-C). On the basis of these synteny relationships, adoption of the new cowpea chromosome numbering for adzuki bean and mung bean would be straightforward. This would facilitate reciprocal exchange of genomic information on target traits from one *Vigna* species to another.

**Figure 3.**
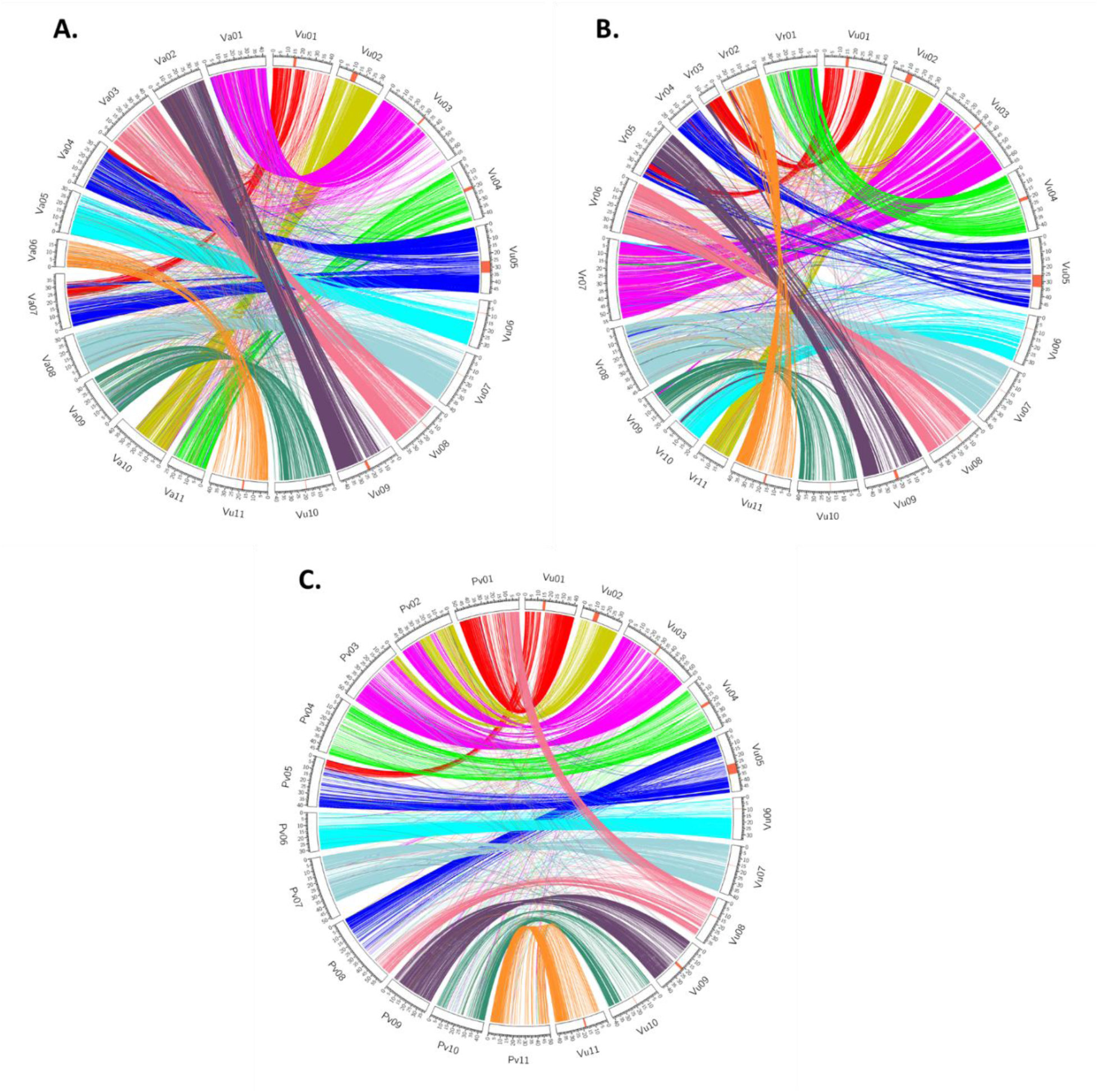
Synteny view between cowpea (Vu; *Vigna unguiculata*) and other closely related diploid species including (A) adzuki bean (Va*; Vigna angularis*), (B) mung bean (Vr; *Vigna radiata*), and common bean (Pv; *Phaseolus vulgaris*) using the new cowpea chromosome numbering system.

### Repetitive elements and genome expansion

Using the same computational pipeline as for *V. unguiculata* (Vu), the repeats of the *V. angularis*^25^ (Va) and *V. radiata*^39^ (Vr) genomes also were annotated. Previous analyses placed cowpea phylogenetically closer to mung bean (Vr) than to adzuki bean (Va)^40^, although the Va and Vr genomes are relatively similar in size, with cowpea respectively 11% and 12% larger. The annotated repeat spaces in the three genomes were examined to make inferences on their evolution. Comparing Vu with Vr, 94% of the 56 Mbp size difference can be explained by the differential abundance of TEs, and 57% by the differential abundance of superfamily *Gypsy* retrotransposons alone (Supplementary Table 15). The differential abundance of *Gypsy* elements in cowpea amounts to 58% and 56% of the total contribution of TEs to its genome size difference with mung bean and adzuki bean, respectively. The non-LTR retrotransposons, composed of SINEs and LINEs, appear to have played only a minor role in genome size enlargement in cowpea. Helitrons contributed 10% (vs. Vr) or 11% (vs. Va) to the expansion of the cowpea genome, and increased in genome share by an order of magnitude. The DNA TEs together contributed 38% of the size difference between Vu and Vr and 40% between Vu and Va. CACTA contributed about the same amount (Va), or 35% more (vs. Vr) of DNA as hAT elements, to this growth. For both Vr and Va, far fewer unidentified LTR retrotransposons (RLX) were found than in the Vu genome, perhaps because the Vu genome appears to be less fragmented and more complete than the former two. Expansion of SSR content was very moderate in Vu vs Vr, and comprised a smaller genome share than in Va.

A similar comparison was made to the 473 Mb genome assembly of *Phaseolus vulgaris*^23^ (Pv) with a genome estimated to be only 9% smaller (587 Mbp; http://data.kew.org/cvalues). However, Pv has a higher TE content than cowpea, 45.2% vs. 39%, of which 39% vs. 33% are retrotransposons. In Pv the *Gypsy* elements comprise 25% of the genome vs. 18% in *V. unguiculata*, although the *Copia* elements are 2% less abundant than in cowpea. There are 23.5 Mb more *Gypsy* elements annotated in the *P. vulgaris* assembly than in Vu, although the total TE coverage is only 10.8 Mb greater in Pv than in cowpea. While the assemblies represent similar shares of the estimated genomes (Vu, 81.1%; Pv, 80.5%), the contig N50 for *P. vulgaris* is 0.395 Mb vs. 10.9 Mb for Vu. These data may indicate that the true *P. vulgaris* genome is considerably larger than estimated by Feulgen densitometry, with the large fraction of TEs interfering with contig assembly.

Taken together, the cross-species comparisons suggest that differences in genome size in *Vigna* can be largely explained by TE abundance, especially by that of *Gypsy* retrotransposons. This can result from either differential amplification recently, or differential retention of ancient insertions. In the grasses, comparison, e.g., of the *Brachypodium distachyon*^42^ and *Hordeum vulgare*^43^ genomes suggests that differences in *Gypsy* content are largely due to differential retention. However, among the legumes examined here, annotated full-length retrotransposons appear to be of recent origin (less than 0.5 million years) in *P. vulgaris*^23^.

### Gene family changes in cowpea

To identify genes that have significantly increased or decreased in copy number in cowpea, we analyzed a set of 18,543 families from the Legume Information System (https://legumeinfo.org/search/phylotree and https://legumeinfo.org/data/public/Gene_families/), constructed to capture gene sets originating at the legume taxonomic depth, based on orthology relationships and per-species synonymous-site rates for legume species and outgroup species. These families include 14 legume species – six of which are from the Phaseoleae tribe (soybean, common bean, adzuki bean, mung bean, pigeon pea, and cowpea - from this study). Among the 185 gene families in the top percentile in terms of cowpea gene membership in the family relative to average membership per legume species, the families include several in the following superfamily groups: NBS-LRR disease resistance genes, various receptor-like protein kinases, defensins, ribosomal proteins, NADH-quinone oxidoreductase components (Supplementary Table 16). All of these families have been observed to occur in large genomic arrays, which can expand or contract, likely through slipped-strand mispairing of paralogous genes^44–46^.

Gene families lacking cowpea membership are more difficult to interpret biologically, as these tend to be smaller gene families, likely showing stochastic effects of small families “falling out of” larger superfamilies, due to extinction of clusters of genes or to artifactual effects of family construction. Among 18,543 legume gene families were 2,520 families without cowpea gene membership, which is comparable to the average number of families without membership (3,057) for six other sequenced genomes in the Phaseoleae. The 2,520 “no-cowpea” families were enriched for the following superfamilies: UDP-glycosyltransferases, subtilisin-like serine proteases, several kinase superfamilies, several probable retrotransposon-related families, FAR1-related proteins, and NBS-LRR disease resistance families (Supplementary Table 16). These large superfamilies are generally organized in large genomic clusters that are subject to expansion and contraction^44, 46, 47^. Several families in cowpea are notable for copy-number differences relative to other sequenced species in *Vigna* (adzuki bean and mung bean). The SAUR-like auxin superfamily contains 138 annotated genes in cowpea, vs. 90 and 52 in adzuki and mung bean, respectively. The NBS-LRR superfamily contains 402 annotated genes, vs. 272 and 86 in adzuki and mung bean, respectively (Supplementary Table 16). In both superfamilies, adzuki and mung bean may have lost gene copies, rather than cowpea gaining genes, or their assemblies underrepresent them due to technological difficulties with short read assemblies capturing such clusters. The cowpea gene counts are more typical of the other annotated Phaseoleae species: 252 and 130 SAUR genes in *Phaseolus* and *Cajanus*, respectively, and 341 and 271 NBS-LRR genes in *Phaseolus* and *Cajanus*, respectively (Supplementary Table 16).

### Identification of a candidate gene for multiple organ gigantism

Crop domestication typically involved size increases of specific organs harvested by humans^48^. Recently, a genomic region related to increased organ size in cowpea was identified on Vu08 using a RIL population derived from a domesticated × wild cross^9^. This region contains a cluster of QTLs for pod length, seed size, leaf length, and leaf width (*CPodl8*, *CSw8*, *CLl8*, *CLw8*)^9^. The reference genome sequence described here was used to further investigate this domestication hotspot, which spans 2.21 Mb and includes 313 genes. Syntenic regions in the common bean genome were identified, the largest of which is located on common bean chromosome 8 (Pv08). That region contains a total of 289 syntelogs, which were then compared with the list of common bean genes associated with domestication available from Schmutz et al.^23^. The intersection of these two lists contained only a single gene, *Phvul.008G285800*, a *P. vulgaris* candidate gene for increased seed size that corresponds to cowpea *Vigun08g217000*. This gene codes for a histidine kinase 2 that is expressed in several cowpea tissues including root, seed, pod and leaf (https://legumeinfo.org). The Arabidopsis ortholog *AHK2* (*AT5G35750.1*) is a cytokinin receptor that has been shown to regulate, among other things, plant organ size^49, 50^. All this evidence suggests that *Vigun08g217000* is a candidate gene for further investigation.

## ONLINE METHODS

### Estimation of genome size

Flow cytometric estimation of genome size followed the protocol of Doležel et al.^12^. Briefly, suspensions of cell nuclei were prepared from 50 mg of young leaf tissue of *V. unguiculata* IT97K-499-35, and of *S. lycopersicum* cv. Stupické polní rané (2C = 1.96 pg DNA) as internal standard. The tissues were chopped using a razor blade in 0.5 ml Otto I solution in a glass Petri dish. The homogenate was filtered through a 50 μm nylon mesh to remove debris and kept on ice until analysis. Then, 1 ml Otto II solution containing 50 μg ml–^1^ propidium iodide and 50 μg ml–^1^ RNase was added and the sample was analyzed by CyFlow Space flow cytometer (Sysmex Partec, Görlitz, Germany). The threshold on the PI detector was set to channel 40 and no other gating strategy was applied. Five thousand events were acquired in each measurement. The resulting histograms of relative DNA content (Supplementary Figure 12) comprised two major peaks representing nuclei in G1 phase of cell cycle. The ratio of G1 peak positions was used to calculate DNA amount of *V. unguiculata*. Five different plants of IT97K-499-35 were analyzed, each of them three times on three different days and the mean 2C DNA amount of *V. unguiculata* was calculated. Genome size was determined using the conversion factor 1 pg = 0.978 Mbp^51^. The values were then considered relative to previous estimates by Parida et al.^13^ and Arumuganathan and Earle^4^. The former estimate was obtained using Feulgen microdensitometry, which is considered a reliable method, and perfect agreement has been observed between flow cytometric and microspectrophotometric estimates^52^. Also, the amount of nuclear DNA assigned to *Allium cepa* (2C = 33.5 pg), which Parida et al.^13^ used as the reference standard differed only by 1.39 pg (4.1 %) from 34.89 pg assigned to *A. cepa* by Doležel et al.^12^. Thus, the higher estimate of DNA amount by Parida et al.^13^ could be due to incomplete removal of formaldehyde fixative prior to staining with Schiff’s reagent, which binds to free aldehyde groups^53^. On the other hand, the present estimate and that of Arumuganathan and Earle.^4^ were obtained using flow cytometry and using a similar protocol. The small difference between genome size estimates for *V. unguiculata* could be due to different values assigned to reference standards. While Arumuganathan and Earle^4^ used domestic chicken considering 2C of 2.33 pg DNA, the present study used *Solanum lycopersicum* (2C = 1.96 pg/2C) calibrated against human with assigned value of 7.0 pg DNA/2C^54^. If this DNA amount is considered for human, then domestic chicken has 2C=2.5 pg DNA^55^ and the estimate by Arumuganathan and Earle^4^ would increase to 658 Mb. However, other factors may be responsible for the observed difference, such as instrument variation between laboratories^52^ and actual differences between the accessions analyzed.

To estimate the cowpea genome size using the k-mer distribution, we processed 168M 149-bp paired-end Illumina reads for a total of about 50 billion bp. Supplementary Figure 12 shows the frequency distribution of 27-mers produced with KAT (https://github.com/TGAC/KAT). The x-axis represents the 27-mer multiplicity, the y-axis represents the number of 27-mers with that multiplicity. The peak of the distribution is 56, which represents the effective coverage. The total number of 27-mers in the range x=2-10000 is 31.381×10^9^. As it is usually done, 27-mers that appear only once are excluded because they are considered erronous, i.e., to contain sequencing errors. The estimated genome size is thus 31.381×10^9^/56 = 560,379,733bp.

### Bionano Genomics optical maps

High molecular weight (HMW) cowpea DNA was isolated by Amplicon Express (Pullman, WA) from nuclei purified from young etiolated leaves (grown in the dark) of 100% homozygous, pure seeds of IT97K-499-35. The material was screened for homozygosity by genotyping with the Cowpea iSelect Consortium Array^5^ (Supplementary Table 17). The nicking endonucleases Nt.*BspQ*I and Nb.*BssS*I (New England BioLabs, Ipswich, MA) were chosen to label DNA molecules at specific sequence motifs. The nicked DNA molecules were then stained according to the instructions of the IrysPrep Reagent Kit (Bionano Genomics, San Diego, CA), as per Luo et al.^56^. The DNA sample was loaded onto the nano-channel array of an IrysChip (Bionano Genomics, San Diego, CA) and then automatically imaged using the Irys system (Bionano Genomics, San Diego, CA). For the *BspQ*I map, seven separate runs (132 unique scans) were generated, and a total of 108 Gb (~170x genome equivalent) of raw DNA molecules (>100 kb) were collected. Molecules of at least 180 kb in length were selected to generate a BNG map assembly. Supplementary Table 1 shows the summary of raw molecule status and the BNG *BspQ*I map assembly. For the *BssS*I map, five separate runs (123 unique scans) were generated, and a total of 186 Gb (~310x genome equivalent) of DNA raw molecules (> 20 kb; 133 Gb molecules > 100 kb) were collected. Molecules of at least 180 kb in length were selected to generate a BNG map assembly. Supplementary Table 2 shows the summary of raw molecules status and the BNG *BssS*I map assembly.

### Whole-genome shotgun sequencing and assembly

#### High molecular weight gDNA and library preparation

Pure seeds of the fully inbred cowpea accession IT97K-499-35 were sterilized and germinated in the dark in crystallization dishes with filter paper and a solution containing antibacterial (cefotaxime, 50 µg/ml) and antifungal (nystatin, 100 units/ml) agents. About 70 g of seedling tissue was collected, frozen in liquid nitrogen, stored at −80C and shipped on dry ice. High molecular weight gDNA was prepared from nuclei isolated from the seedling tissue by Amplicon Express (Pullman, WA).

#### Pacific Biosciences sequencing

Pacific Biosciences reads were generated at Washington State University (Pullman, WA) following the “Procedure and Checklist-20 kb Template Preparation Using BluePippin Size Selection System” (P/N 100-286-000-5) protocol provided by Pacific Biosciences (Menlo Park, CA) and the Pacific Biosciences SMRTbell Template Prep kit 1.0 (P/N 100-259-100). Resulting SMRTbell libraries were size selected using the BluePippin (Sage Biosciences) according the Blue Pippin User Manual and Quick Guide. The cutoff limit was set to 15-50 kb to select SMRTbell library molecules with an average size of 20 kb or larger. The Pacific Biosciences Binding and Annealing calculator determined the appropriate concentrations for the annealing and binding of the SMRTbell libraries. SMRTbell libraries were annealed and bound to the P6 DNA polymerase for sequencing using the DNA/Polymerase Binding Kit P6 v2.0 (P/N100-372-700). The only deviation from standard protocol was to increase the binding time to 1-3 hours, compared to the suggested 30 minutes. Bound SMRTbell libraries were loaded onto the SMRT cells using the standard MagBead protocol, and the MagBead Buffer Kit v2.0 (P/N 100—642-800). The standard MagBead sequencing protocol followed the DNA Sequencing Kit 4.0 v2 (P/N 100-612-400) which is known as P6/C4 chemistry. PacBio RS II sequencing data were collected in six-hour movies and Stage Start was enabled to capture the longest sub-reads possible.

#### Sequence quality control

First, CLARK and CLARK-*S*^57^ were used to identify possible contamination from unknown organisms. CLARK and CLARK-*S* are classification tools that use discriminative (spaced, in the case CLARK-S) *k*-mers to quickly determine the most likely origin of each input sequence (*k=21* and *k=31*). The target database for CLARK/CLARK-*S* was comprised of: (i) a representative sample of ~5,000 bacterial/viral genomes from NCBI RefSeq, (ii) human genome, *Homo sapiens*, assembly GRCh38, (iii) Illumina-based cowpea draft genome, *Vigna unguiculata*^5^, assembly v0.03), (iv) soybean, *Glycine max*^30^, assembly Gmax_275_v2.0, (v) common bean, *Phaseolus vulgaris*^23^, assembly Pvulgaris_218_v1.0, (vi) adzuki bean, *Vigna angularis*^25^, assembly adzuki.ver3.ref.fa.cor, (vii) mung bean, *Vigna radiata*^39^, assembly Vradi.ver6.cor, and (viii) a nematode that attacks the roots of cowpea, *Meloidogyne incognita*^58^, assembly GCA_900182535.1_Meloidogyne_incognita_V3.

#### Whole genome assemblies

Eight draft assemblies were generated, six of which were produced with CANU v1.3^16, 17^, one with Falcon v0.7.3^18^, and one with Abruijn v0.4^19^. Hinge v0.41^59^ was also tested on this dataset, but at that time the tool required the entire alignment file (over 2Tb) to fit in primary memory and we did not have the computational resources to handle it. CANU v1.3 was run with different settings for the error correction stage on the entire dataset of ~6M reads (two CANU runs were optimized for highly repetitive genomes). Falcon and Abruijn were run on 3.54 M error-corrected reads produced by CANU (30.62Gbp, or 49.4x genome equivalent). Each assembly took about 4-15 days on a 512-core Torque/PBS server hosted at UC Riverside.

#### Removal of contaminants from the assemblies

To remove “contaminated” contigs, two sets of reference genomes were created, termed the *white* list and the *black* list. Black-list genomes included possible contaminants, whereas white-listed genomes included organisms evolutionarily close to cowpea. The black list included: (i) *Caulobacter segnis* (NCBI accession GCF 000092285.1), (ii) *Rhizobium vignae* (NCBI accession GCF 000732195.1), (iii) *Mesorhizobium sp. NBIMC P2-C3* (NCBI accession GCF 000568555.1), (iv) *Streptomyces purpurogeneiscleroticus* (NCBI accession GCF 001280155.1), (v) *Caulobacter vibrioides* (NCBI accession GCF 001449105.1), (vi) mitochondrion of *Vigna radiata*^60^ (NCBI accession NC_015121.1), (vii) mitochondrion of *Vigna angularis* (NCBI accession NC_021092.1), (viii) chloroplast of *Vigna unguiculata* (NCBI accession NC_018051.1 and KJ468104.1), and (ix) human genome (assembly GRCh38). The white list included the genomes of: (i) soybean (*Glycine max*^30^, assembly Gmax_275_v2.0), (ii) common bean (*Phaseolus vulgaris*^23^, assembly Pvulgaris_218_v1.0), (iii) adzuki bean (*Vigna angularis*^25^, assembly adzuki.ver3.ref.fa.cor), (iv) mung bean (*Vigna radiata*^39^, assembly Vradi.ver6.cor), and (v) Illumina-based cowpea draft genome (*Vigna unguiculata*^5^, assembly v.0.03). Each assembled contig was BLASTed against the “white” genome and the “black” genomes, and all high-quality alignments (e-score better than 1e^−47^ corresponding to a bit score of at least 200, and covering at least 10% of the read length) were recorded. The percentage of each contig covered by white and black high-quality alignments was computed by marking each alignment with the corresponding identity score from the output of BLAST. When multiple alignments covered the same location in a contig, only the best identity alignment was considered. The sum of all these identity scores was computed for each contig, both for the black and the white list. These two scores can be interpreted as the weighted coverage of a contig by statistically significant alignments from the respective set of genomes. A contig was considered contaminated when the black score was at least twice as high as the white score. Chimeric contigs were identified by mapping them against the optical maps using RefAligner (Bionano Genomics), then determining at what loci to break chimeric contigs by visually inspecting the alignments using IrysView (Bionano Genomics).

#### Stitching of contaminant-free assemblies and polishing

Our method (i) uses optical map(s) to determine small subsets of assembled contigs from the individual assemblies that are mutually overlapping with high confidence, (ii) computes a minimum tiling path (MTP) of contigs using the coordinates of the contigs relative to the optical map, and (iii) attempts to stitch overlapping contigs in the MTP based on the coordinates of the contigs relative to the optical map. A series of checks are carried out before and after the stitching to minimize the possibility of creating mis-joins. Additional details about the stitching method can be found in Pan et al.^20^. The final stitched assembly was then polished via the PacBio Quiver pipeline (RS_resequencing.1 protocol) in SMRT Portal v2.3.0 (Patch 5) by mapping all the PacBio subreads against the assembly. The polishing step took about seven days on a 40-core server at UC Riverside.

#### Scaffolding via optical maps

Scaffolds were obtained from the polished assembly via the Kansas State University (KSU) stitching pipeline^21^ in multiple rounds. A tool called XMView (https://github.com/ucrbioinfo/XMView) developed in-house that allows the visual inspection of the alignments of assembled contigs to both optical maps simultaneously was used to identify chimeric optical molecules that had to be excluded from the scaffolding step. The KSU stitching pipeline was iterated four times, alternating *BspQ*I and *BssS*I (twice each map).

#### Pseudo-molecule construction via anchoring to genetic maps

Pseudo-molecules were obtained by mapping the SNP markers in ten genetic maps (Supplementary Table 4) to the sequence of the scaffolds. Seven of these genetic maps were previously generated, five of which are available from Muñoz-Amatriaín et al.^5^ and one each from Santos et al.^27^ and Lo et al.^9^ The remaining three genetic maps were generated as part of this study by genotyping three additional RIL (Recombinant Inbred Line) populations with the Cowpea iSelect Consortium Array^5^. SNP calling and curation were done as described by Muñoz-Amatriaín et al.^5^, and linkage mapping was performed using MSTmap^61^. Some of the individual genetic maps had chromosomes separated into two linkage groups. In those cases, the most recent cowpea consensus genetic map^5^ was used to join them by estimating the size of the gap (in cM). The locations of SNPs on the scaffolds was determined by BLAST using the 121 bp SNP design sequences, filtering for alignments with an e-score of 1e^−50^ or better. The final ordering and orientation of the scaffold was produced by ALLMAPS^22^ from the SNP locations corresponding to the ten genetic maps. As noted elsewhere, 46 Mb of assembled sequences were not anchored. In addition, 24.5 Mb of the anchored sequences were oriented arbitrarily.

#### Annotation method and estimation of centromere positions

Transcript assemblies were made from ~1.5 B pairs of 2×100 paired-end Illumina RNAseq reads^11, 27^ using PERTRAN (Shu, unpublished). 89,300 transcript assemblies were constructed using PASA^62^ from EST-derived UNIGENE sequences (P12_UNIGENES.fa^63^; harvest.ucr.edu) and these RNAseq transcript assemblies. Loci were determined by transcript assembly alignments and/or EXONERATE alignments of proteins from Arabidopsis (*Arabidopsis thaliana*), common bean, soybean, medicago, poplar, rice, grape and Swiss-Prot proteomes to repeat-soft-masked *Vigna unguiculata* genome using RepeatMasker^64^ with up to 2 kb extension on both ends unless extending into another locus on the same strand. The repeat library consisted of de novo repeats identified by RepeatModeler^65^ and Fabaceae repeats in RepBase. Gene models were predicted by homology-based predictors, FGENESH+^66^, FGENESH_EST (similar to FGENESH+, EST as splice site and intron input instead of protein/translated ORF), and GenomeScan^67^, PASA assembly ORFs (in-house homology constrained ORF finder) and from AUGUSTUS via BRAKER1^68^. The best scored predictions for each locus were selected using multiple positive factors including EST and protein support, and one negative factor: overlap with repeats. The selected gene predictions were improved by PASA. Improvement includes adding UTRs, splicing correction, and adding alternative transcripts. PASA-improved gene model proteins were subject to protein homology analysis to the proteomes mentioned above to obtain Cscore and protein coverage. Cscore is a protein BLASTP score ratio to MBH (mutual best hit) BLASTP score and protein coverage is highest percentage of protein aligned to the best of homologs. PASA-improved transcripts were selected based on Cscore, protein coverage, EST coverage, and its CDS overlapping with repeats. The transcripts were selected if the Cscore was larger than or equal to 0.5 and protein coverage larger than or equal to 0.5, or if it had EST coverage, but its CDS overlapping with repeats was less than 20%. For gene models whose CDS overlaps with repeats for more than 20%, its Cscore had to be at least 0.9 and homology coverage at least 70% to be selected. The selected gene models were subjected to Pfam analysis, and gene models whose protein was more than 30% in Pfam TE domains were removed.

The centromere-abundant 455-bp tandem repeat available from Iwata-Otsubo et al.^26^ was BLASTed against cowpea pseudomolecules to identify the approximate start and end positions of cowpea centromeres. Only alignments with an e-score ≤ 1e^−50^ were considered. The chromosome region extending from the beginning of the first hit to the end of the last hit was considered to define the centromeric region of each cowpea chromosome.

### Recombination rate

A polynomial curve fit of cM position as a function of pseudomolecule coordinate was generated using R for each of the eleven linkage groups from each of ten biparental RIL populations. The built-in R function `lm`() was used to compute the linear regression, function `predict`() was used to create the raster objects and function `polynomial`() yielded the polynomial coefficients. For each curve, the best fit from polynomials ranging from 4th to 8th order was selected. The first derivative was then calculated for each of the 110 selected polynomials to represent the rate of recombination as cM/nucleotide. The mean values of the recombination rates (first derivative values) were then calculated along each of the eleven linkage groups after setting all negative values to zero and truncating values at the ends of each linkage group where the polynomial curve clearly no longer was a good fit. A polynomial was then derived for the mean values along each chromosome/pseudomolecule to represent recombination rate as a function of nucleotide coordinate. Supplementary Table 8 provides the polynomial formulae for each chromosome/pseudomolecule.

### Repeat Analysis

Repeats in the contigs and pseudochromosomes were analyzed using RepeatMasker. An initial library of elements was built by combining the output from Repet, RepeatModeler, LTRharvest/LTRdigest (genometools.org), elements in the Fabaceae section of the RepBase transposon library^69^ and our own custom pipeline. Subsequent *Vigna*-specific libraries were built by iterative searches. The resulting *Vigna*-specific libraries were used again in iterative searches to build the set of elements in the genome. The set was supplemented with elements identified by similarities to expected domains, including LINE integrases for the LINEs and transposases for the DNA transposons. The set was supplemented by searches based on structural criteria typical of various groups of transposable elements. For classifying the repeats, an identity of at least 80% and minimal hit length 80 bp were required.

For the LTR retrotransposons, full-length versions were identified with LTRharvest^70^ using the following parameter settings: “overlaps best -seed 30 -minlenltr 100 -maxlenltr 3000 - mindistltr 100 -maxdistltr 15000 -similar 80 -mintsd 4 -maxtsd 20 -motif tgca -motifmis 1 -vic 60-xdrop 5 -mat 2 -mis −2 -ins −3 -del −3”. All candidates were annotated for PfamA domains with hmmer3 software^71^ and stringently filtered for false positives by several criteria, the main ones being the presence of at least one typical retrotransposon domain (e.g., reverse transcriptase, RNaseH, integrase, Gag) and a tandem repeat content below 5%.

### Identification of genetic variation

Nearly 1M SNPs with strong support were discovered previously by aligning WGS data from 36 diverse accessions to a draft assembly of IT97K-499-35^5^. To position those SNPs on the cowpea reference genome, the 121-base sequences comprised of the SNP position and 60 bases on each side were BLASTed against the cowpea genome assembly with an e-score cutoff of 1e^−50^. Only the top hit for each query was kept. The exact SNP position was then calculated. SNPs previously identified as organellar were excluded, together with those hitting multiple locations in the reference genome sequence.

For detection of insertions and deletions, WGS data from 36 diverse accessions^5^ were used. Reads from each cowpea accession were mapped to the genome assembly using BWA-MEM version 0.7.5a^72^. Variant calling was carried on each resulting alignment using BreakDancer version 1.4.5^31^, with a minimum mapping quality score of 30 and 10 as the minimum number of pair-end reads to establish a connection. The maximum structural variation size to be called by BreakDancer was set to 70 kb. A deletion was considered validated when at least 75% of the SNPs contained in the deletion region were “No Call”.

To validate the inversion, the sequence assembly of the reference genome was compared to that of a cowpea accession typical of California breeding lines via MUMmer^73^, using a minimum exact match of 100 bp and a minimum alignment length of 1 kb. PCR amplifications of the breakpoint regions were performed to further validate the Vu03 inversion. Four accessions were tested for each of the two orientations (type A and type B); these were parental lines of some of the ten genetic maps used for anchoring (Supplementary Figure 9) and included one wild cowpea (TVNu-1158). Two primer pairs were designed for each breakpoint region: one to amplify the reference orientation and another to amplify the opposite orientation (Supplementary Table 17). For the latter, the sequence assembly of the California accession was used to design primers. When primers were designed to amplify the reference orientation, they worked well in type A accessions, but they did not work for the type B accessions (Supplementary Figure 10). When primers were designed to amplify the opposite orientation, there was PCR product only in the type B accessions (Supplementary Figure 10). Only the wild cowpea accession did not yield an amplification product for either of the breakpoints, possibly due to sequence variation within the breakpoint regions.

### Synteny between cowpea and *Phaseolus vulgaris, Vigna radiata* and *Vigna angularis*

The cowpea IT87K-499-35 genome sequence assembly was compared to that of common bean v2.1^23^, adzuki bean^74^ and mung bean^39^ using MUMmer software package v3.23^73^. Alignments were generated using pipeline ‘nucmer’, with a minimum length of an exact match set to 100 bp. Alignments with a length less than 1 kb were filtered out. The output alignments between genomes were visualized using Circos v0.69-3^75^.

### Gene Families

To compare annotated genes in cowpea with those from other legume proteomes, we utilized the legume-focused gene families from the NSF Legume Federation project (NSF DBI#1444806). This is a set of 18,543 gene families, constructed to be monophyletic for the legume family, and including proteomes from cowpea (this study), thirteen other major crop and model legumes, and five non-legume species used for phylogenetic rooting and evolutionary context (Supplementary Table 18). Gene families were generated as follows (summarizing method details from https://github.com/LegumeFederation/legfed_gene_families). All-by-all comparisons of all protein sequences were calculated using BLAST^76^, with post-processing filters of 50% query coverage and 60% identity. The top two matches per query were used to generate nucleotide alignments of coding sequences, which were used, in turn, to calculate synonymous (Ks) counts per gene pair. For each species pair, histograms of Ks frequencies were used as the basis for choosing per-species Ks cutoffs for that species pair in the legumes. A list of all-by-all matches, filtered to remove all pairs with Ks values greater than the per-species-pair Ks cutoff, was used for Markov clustering implemented in the MCL program^77^, with inflation parameter 1.2, and relative score values (transformed from Ks values) indicated with the -abc flag. Sequence alignments were then generated for all families using muscle^78^, and hidden Markov models (HMMs) were calculated using the hmmer package^79^. Family membership was evaluated relative to median HMM bitscores for each family, with sequences scoring less than 40% of the median HMM bitscore for the family being removed. The HMMs were then recalculated from families (without low-scoring outliers), and were used as targets for HMM search of all sequences in the proteome sets, including those omitted during the initial Ks filtering. Again, sequences scoring less than 40% of the median HMM bitscore for the family were removed. Prior to calculating phylogenetic trees, the HMM alignments from the resulting family sets were trimmed of non-aligning characters (characters outside the HMM match states). Phylogenies were calculated using RAxML^80^, with model PROTGAMMAAUTO, and rooted using the closest available outgroup species.

### Identification of a syntelog for increased organ size

The identification of QTLs on Vu08 for organ size (*CPodl8*, *CSw8*, *CLl8*, *CLw8*) is described in Lo, et al.^9^. The SNP markers associated with those QTLs span the genomic region Vu08:36035190-38248903, which contains 313 annotated genes. The corresponding syntenic segment in *Phaseolus vulgaris* (Chr08: 57594596-59622008) was determined using the legumeinfo.org instance of the Genome Context Viewer (GCV)^81^. This region contained 289 Phaseolus genes, of which only one (*Phvul.008G285800*) was present in the intersection with a list of genes associated with domestication (reported in Schmutz, et al.^23^) as determined using functions of cowpeamine and legumemine (https://mines.legumeinfo.org), instances of the InterMine data warehousing system^82^. The cowpea syntelog of that gene is *Vigun08g217000*, according to the genomic segment alignment provided by the GCV using the gene family assignments described above.

## DATA AVAILABILITY

Raw PacBio reads for cowpea accession IT97K-499-35 are available at NCBI SRA sample SRS3721827 (study SRP159026). The genome assembly of cowpea IT97K-499-35 is available through NCBI SRA BioSample accession SAMN06674009 and Phytozome (phytozome.jgi.doe.gov). RNA-Seq raw reads are available as NCBI SRA biosample accessions SAMN071606186 through SAMN071606198, SAMN07194302 through SAMN07194309 and SAMN07194882 through SAMN07194909 and were described in Yao et al.^11^ and Santos et al.^27^. EST sequences and their GenBank accession numbers are available through the software HarvEST:Cowpea (harvest.ucr.edu) and were described in Muchero et al.^63^.

## ACKNOWLEDGMENTS

Authors thank: Panruo Wu, Laxmi Buyhan, Zizhong Chen and Tamar Shinar (University of California Riverside, CA) for allowing access to their HPC server; Suresh Iyer, Amy Mraz, Jon Wittendorp, and Robert Bogden (Amplicon Express, Pullman, WA) for DNA extraction and library prep; Derek Pouchnik and Mark Wildung (Washington State University, Pullman, WA) for PacBio sequencing and library preparation; Thiru Ramaraj (National Center for Genome Resources), Matthew Seetin and Christopher Dunn (Pacific Biosciences of California, Inc., Menlo Park, CA), Brian Walenz (University of Maryland, College Park, MD), John Urban (Brown University, Providence, Rhode Island), and Xingtan “Tanger” Zhang for helpful discussion on assembly tools, trouble shooting, usage and parameter selection; Haibao Tang (JVCI) for help with ALLMAPS; Suk-Ha Lee (Seoul National University, South Korea) for allowing access to the most recent mung bean and adzuki bean genome assemblies; David Goodstein (Joint Genome Institute, Walnut Creek, CA) for assistance coordinating genome annotation; Yi-Ning Guo and Savannah St Clair (UC Riverside, CA) for technical assistance with DNA preparation; and Ira Herniter (UC Riverside, CA) for helpful discussion. This work was supported by the NSF IOS-1543963 (“Advancing the Cowpea Genome for Food Security”), NSF IIS-1526742 (“Algorithms for Genome Assembly of Ultra-Deep Sequencing Data”) and NSF IIS-1814359 (“Improving *de novo* Genome Assembly using Optical Maps”). Estimation of the genome size was supported by the Czech Ministry of Education, Youth and Sports (award LO1204 from the National Program of Sustainability I). The analysis of gene families was provided through by in-kind contributions from the USDA Agricultural Research Service, project 5030-21000-069-00-D, while the repetitive elements analysis was supported by the Academy of Finland “Papugeno” (Decision 298314).

## AUTHOR CONTRIBUTIONS

TJC, StL and MMA conceived and supervised the study. StL coordinated the sequencing and executed the assembly with help from SIW. RO identified contamination sequences. TZ and MCL generated the optical maps. MMA, SaL and AN generated genetic maps. JV, PAR and JS contributed to the generation of transcriptome data. SS generated gene annotations. AHS and JT annotated and analyzed repeats. SIW, MMA and TJC contributed to SNP annotation and analysis. HA and AMH identified structural variants. MMA analyzed and validated the chromosomal inversion with help from SaL, SIW and ADF. TJC and SIW estimated recombination rates. QL performed synteny analyses, identified cowpea centromeres and generated a visualization of the distribution of genes, repeats and genetic variation across the genome. TJC, StL, QL and MMA developed the new chromosome numbering for cowpea. SBC and ADF performed gene family analyses. SaL, SAH and ADF identified the syntelog for multiple organ gigantism. JD and JV estimated the genome size. StL and MMA wrote the manuscript with inputs from TJC, SBC, ADF, JD and AHS.

