## Supplementary_Tables_and_Figures for "The genome of cowpea (*Vigna unguiculata* [L.] Walp.)"

**Supplementary Figure 1.** Distribution of occurrences of 27-mers in 168M 149-bp paired-end reads produced with KAT (https://github.com/TGAC/KAT). The total number of base pairs is about $50x10^9$. The x-axis represents the 27-mer multiplicity, the y-axis represents the number of 27-mers with that multiplicity. The peak of the distribution is 56, which represents the effective coverage. The total number of 27-mers in the range x=2-10000 is $31.381x10^9$ bp (27-mers that appear only once are considered erronous, i.e., to contain sequencing errors). The estimated genome size is thus $31.381x10^9/56$ = 560,379,733bp.

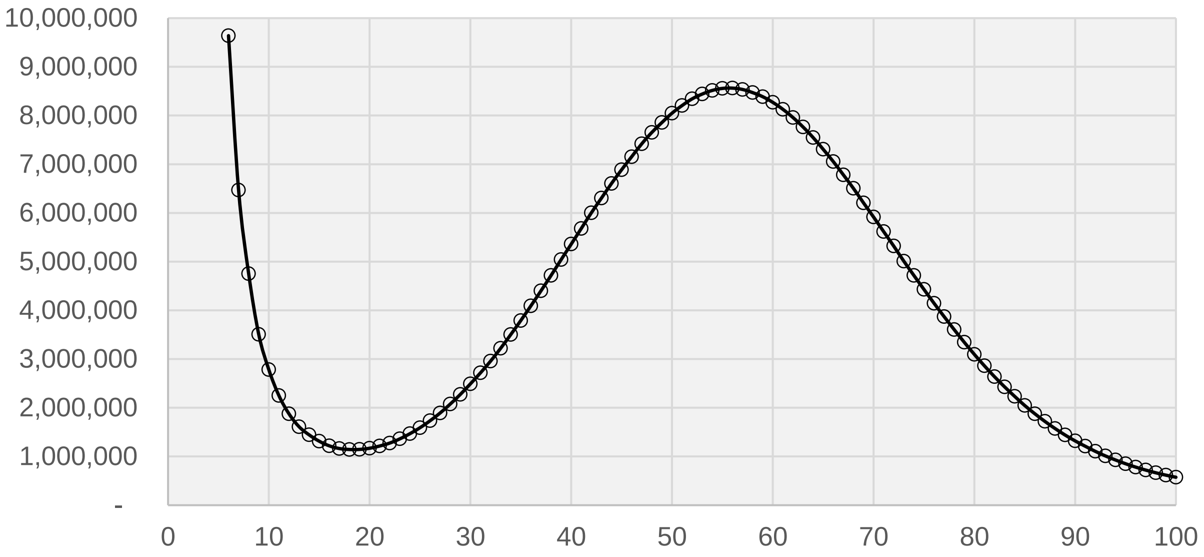

**Supplementary Figure 2.** Pre-filter PacBio read length distribution.

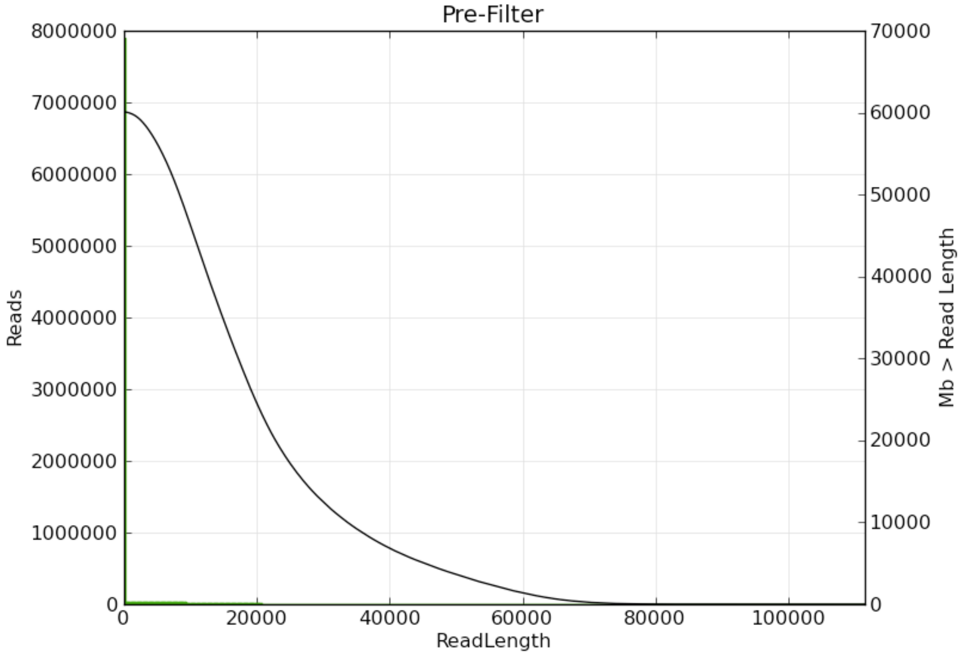

**Supplementary Figure 3.** Post-filter PacBio read length distribution (green).

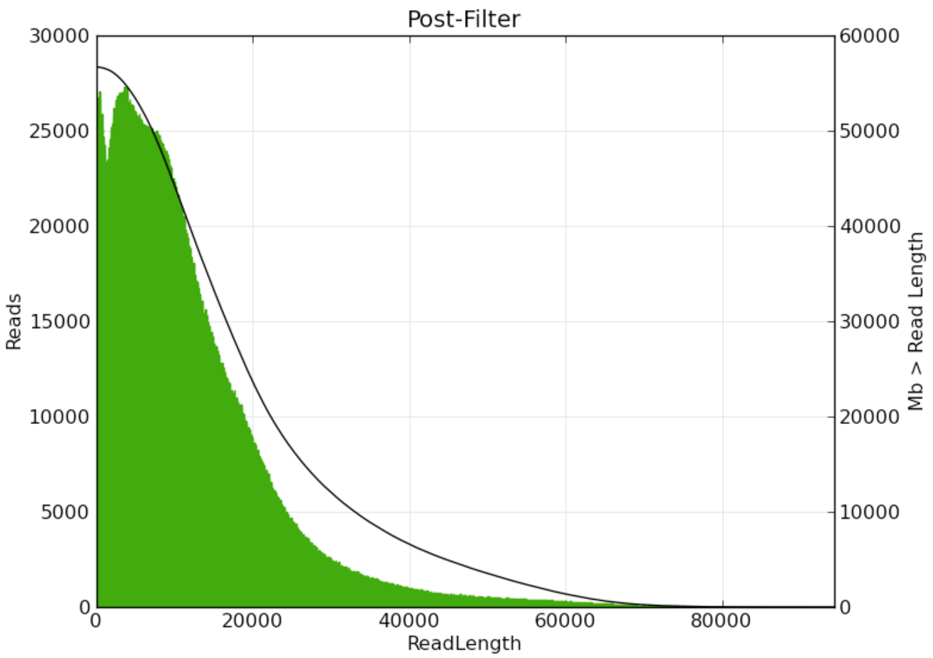

**Supplementary Figure 4.** Subread Filtering PacBio read length distribution (green).

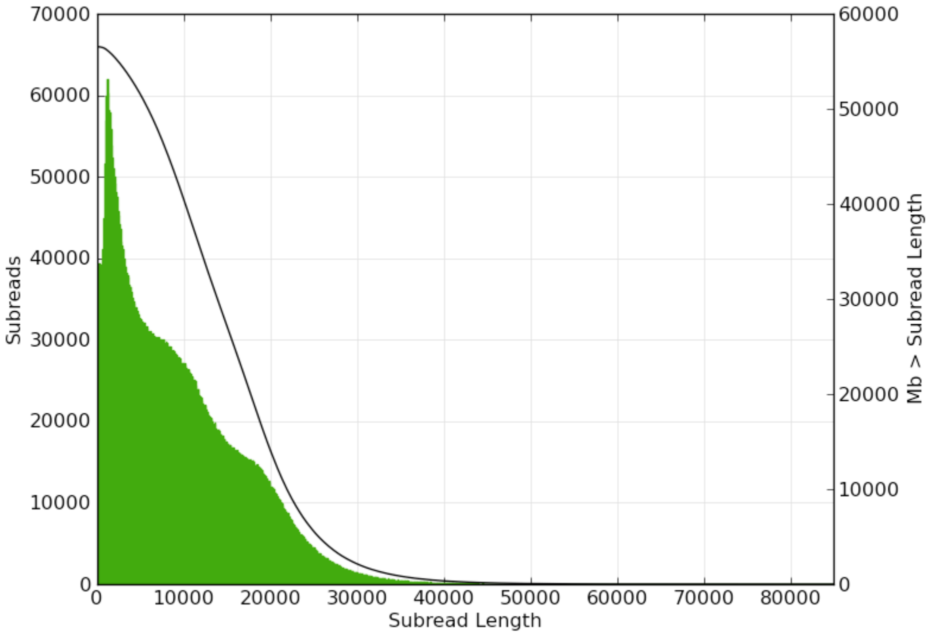

**Supplementary Figure 5.** Post-filter PacBio read quality distribution.

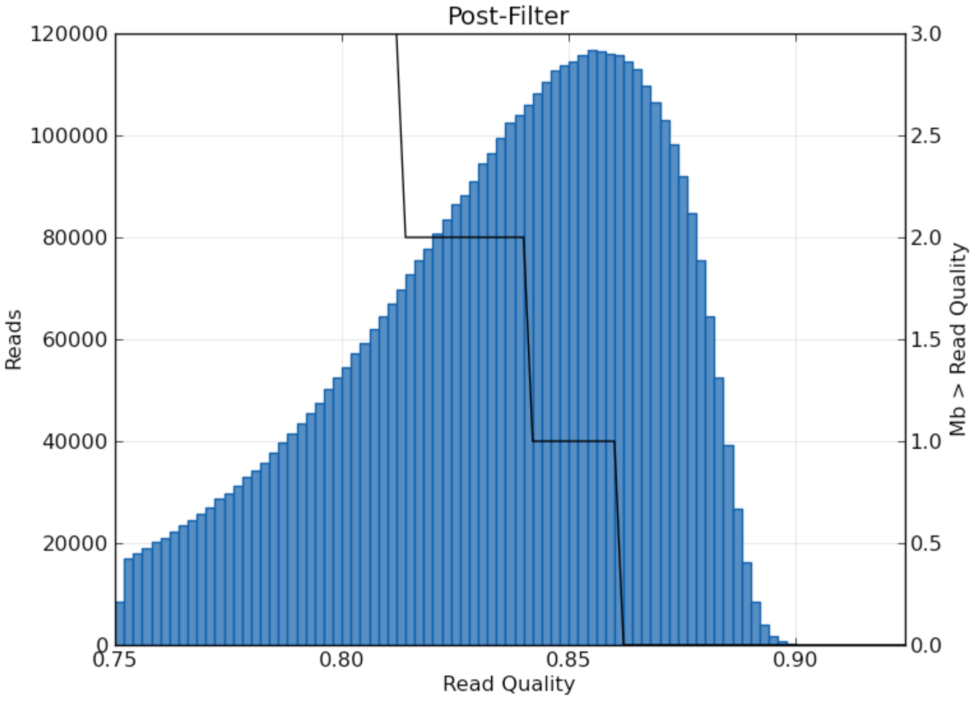

**Supplementary Figure 6.** Synteny view between cowpea (Vu) and common bean (Pv) chromosomes using the previous cowpea chromosome numbering of Muchero et al.[63] and Muñoz-Amatriaín et al.[5]. (A) Circos illustration of synteny. (B) Cowpea chromosomes painted based on syntenic relationships with common bean chromosomes (in different colors). Only syntenic regions with a exact match of 100 bp and a minimum alignment >1kb are colored. Chromosomes indicated with arrows need to be inverted to meet the convention (short arm on top) based on the BAC-FISH analysis of Iwata-Otsubo et al.[26].

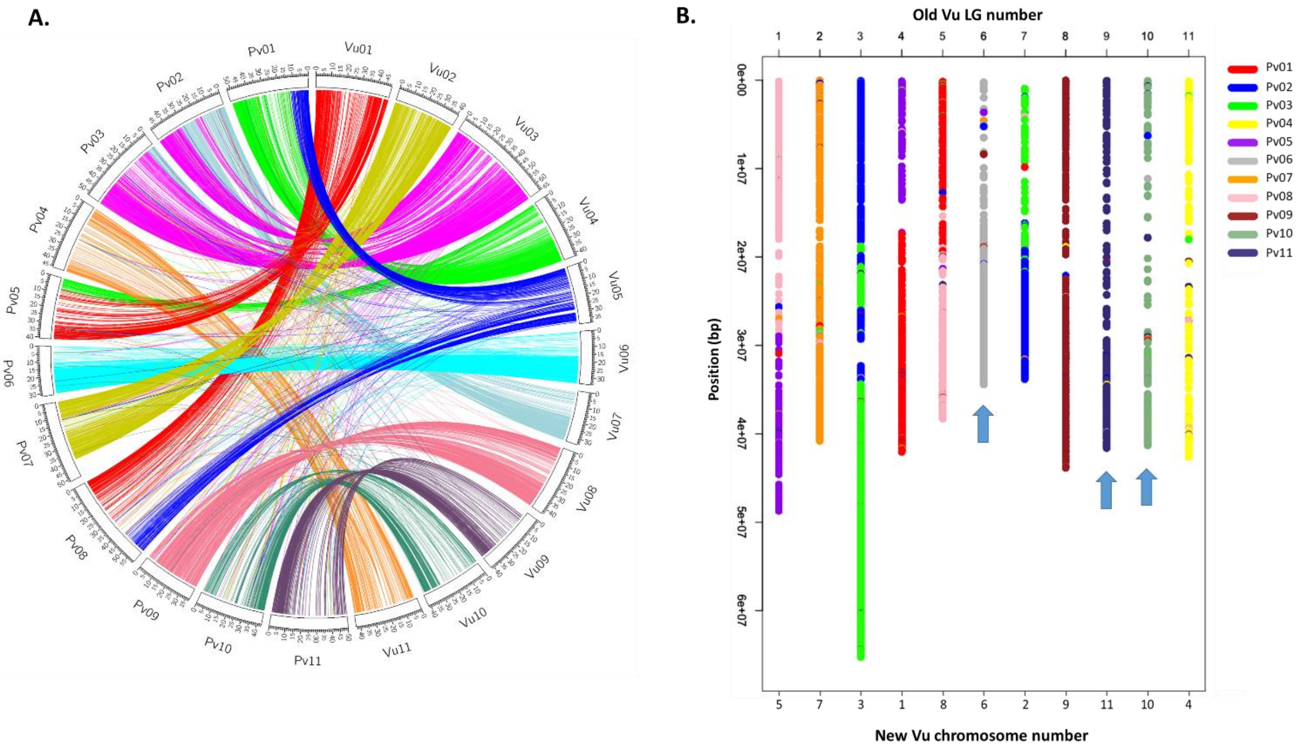

**Supplementary Figure 7.** Comparison between gene density (green line), repeat density (blue line) and recombination rate (red line) across the 11 cowpea chromosomes. Gene and repeat density are measured in 1 Mb non-overlapping windows, while recombination rate is measured in non-overlapping windows of 100 kb. Vertical lines delimit the predicted centromeric regions.

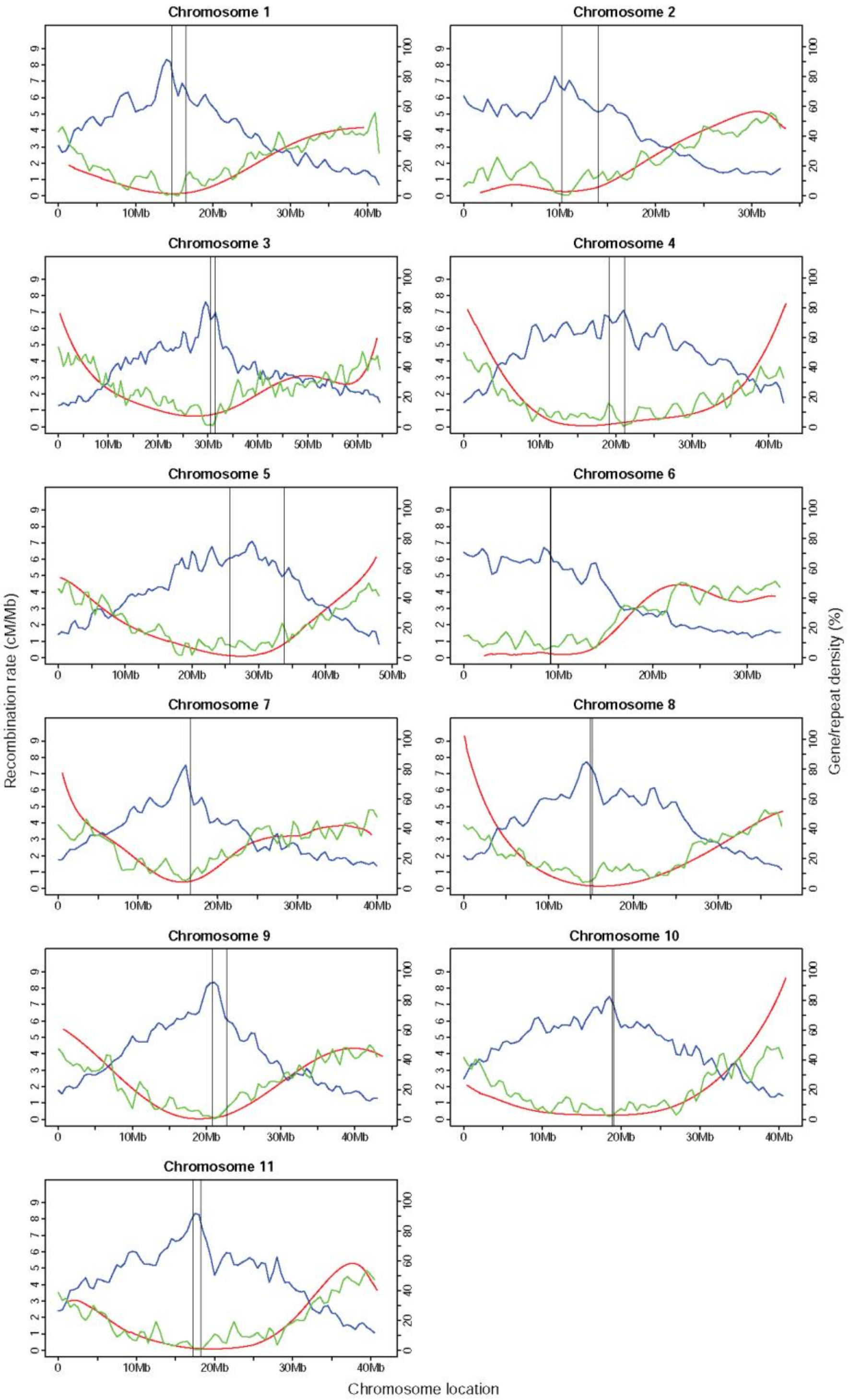

**Supplementary Figure 8.** Chromosome location of SNPs from the "1M list" (in red) and the Illumina iSelect Consortium Array (in blue). Arrows delimit the predicted centromeric regions.

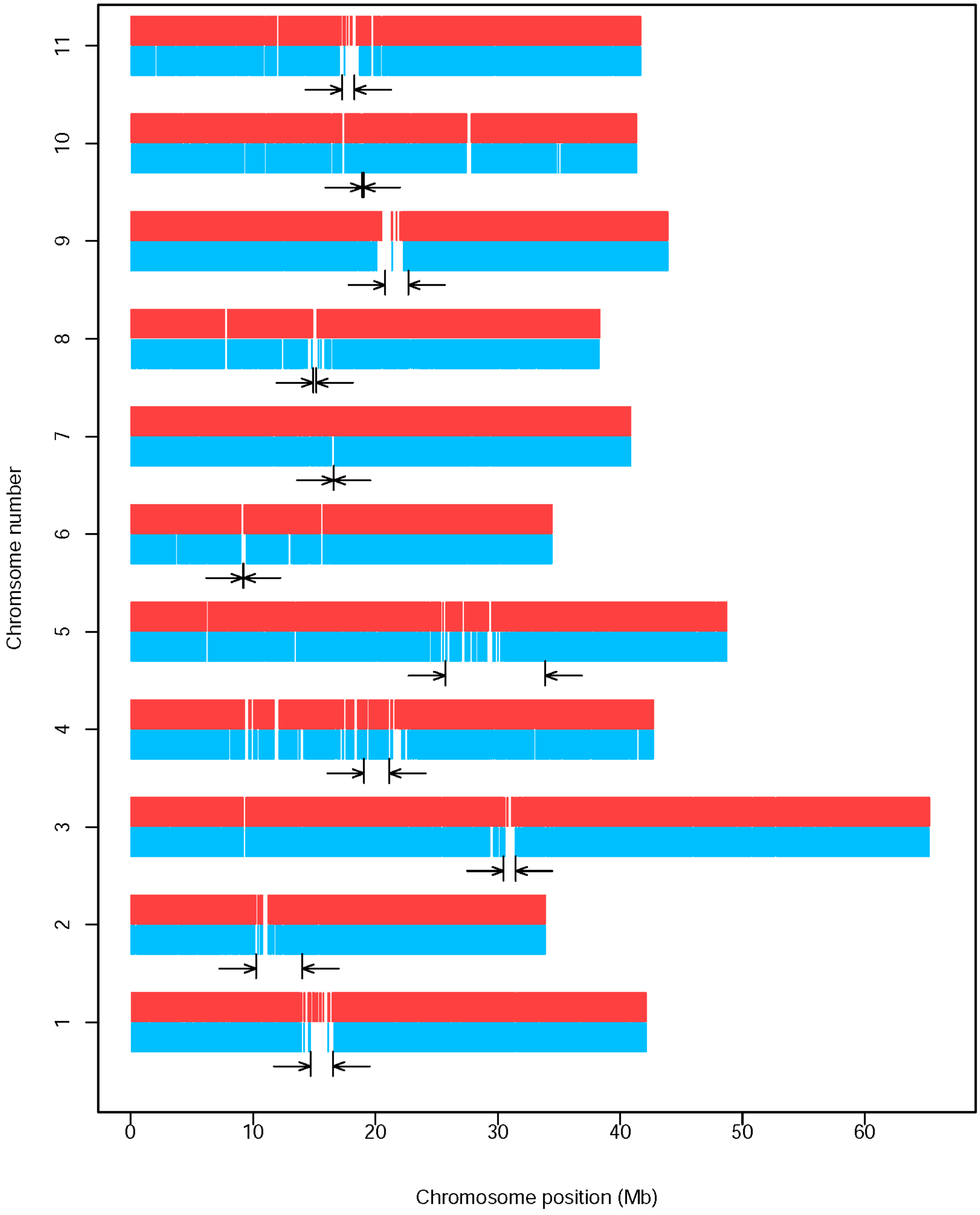

**Supplementary Figure 9.** Cowpea pseudomolecule Vu03 reconstructed from 10 genetic maps using ALLMAPS. (A) Alignments between physical positions on pseudomolecule Vu03 and genetic map positions. (B) Scatter plots of genetic (y-axes) vs. physical (x-axes) map positions. Rho ($\rho$) represents the Pearson correlation coefficient. The chromosomal inversion can be observed in 7 out of the 10 genetic maps (red arrow). 27B=CB27xIT82E-18; 27I=CB27xIT97K-566-6; 27U=CB27xUCR779; 46I=CB46xIT93K-503-1; 5I=524BxIT84S-2049; NF=NullxFN-2-9-04; SV=SanzixVita7; TI=Tvu-14676xIT84S-2246-4; WC=IT99K-573-1-1xTVNu-1158; ZZ=ZN016xZhijiang282.

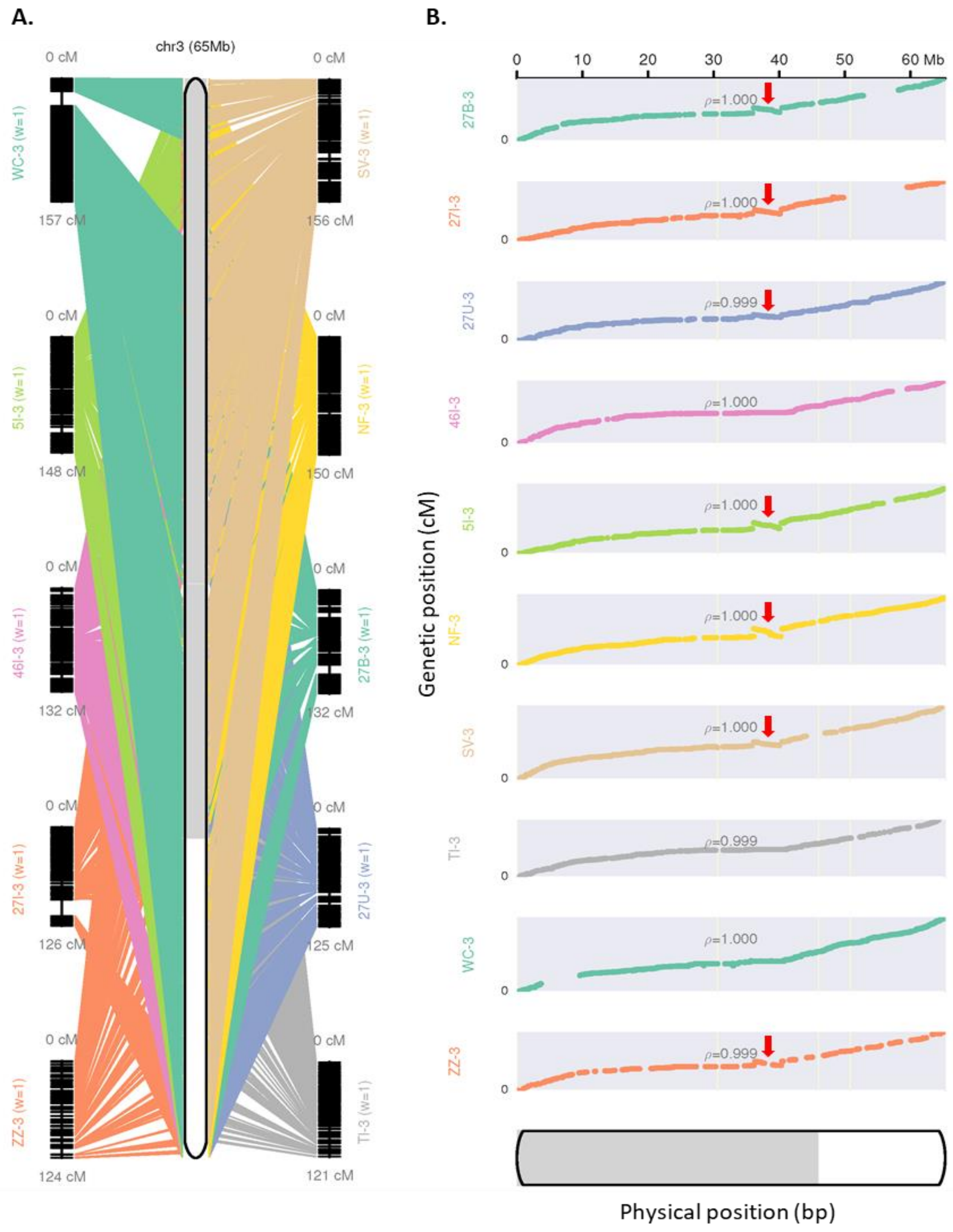

**Supplementary Figure 10.** PCR amplifications of the regions surrounding the two breakpoints of the inversion. A total of four accessions for each of the two orientations were tested. Type A refers to accessions having the same orientation as the reference genome, while type B indicates accessions having the opposite-to-reference orientation. The reference genome sequence (IT97K-499-35) was used for designing primers for the "reference" orientation, while the type B sequence assembly (unpublished) was used to design primers for the opposite orientation.

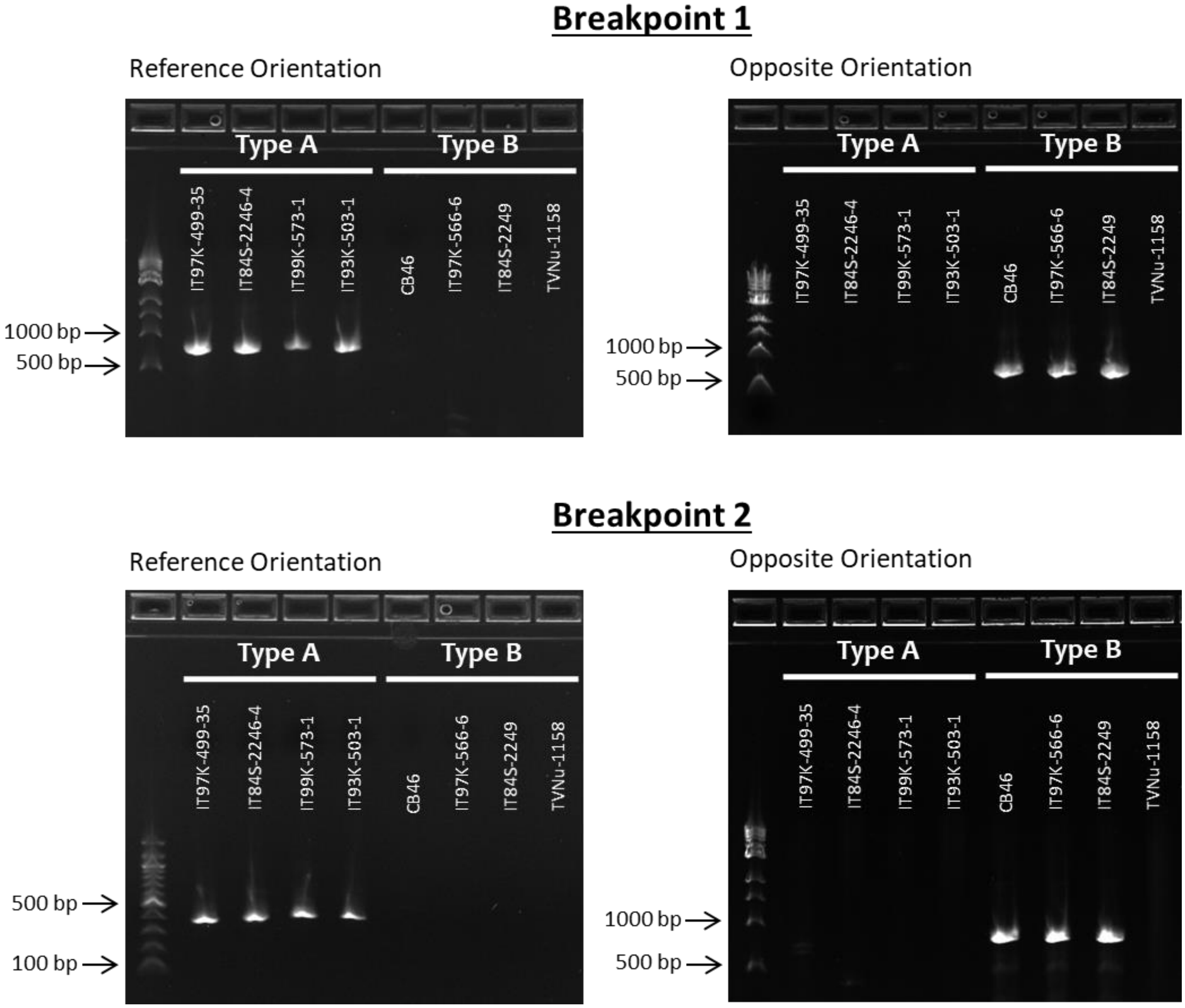

**Supplementary Figure 11.** Comparison between the Vu03 chromosomal inversion region in cowpea (y-axis) and its syntenic region in *Vigna angularis* (vigan.Shumari). Dots in the plot represent gene pairs likely to be orthologous between the two species, as assessed by their assignment to the same gene family (represented by the dot colors) and their presence in these largely co-linear blocks. The break in the middle of the inverted segment (~36-41Mb in the *V. angularis* chromosome), corresponds to a block further translocated with respect to the cowpea region (not shown).

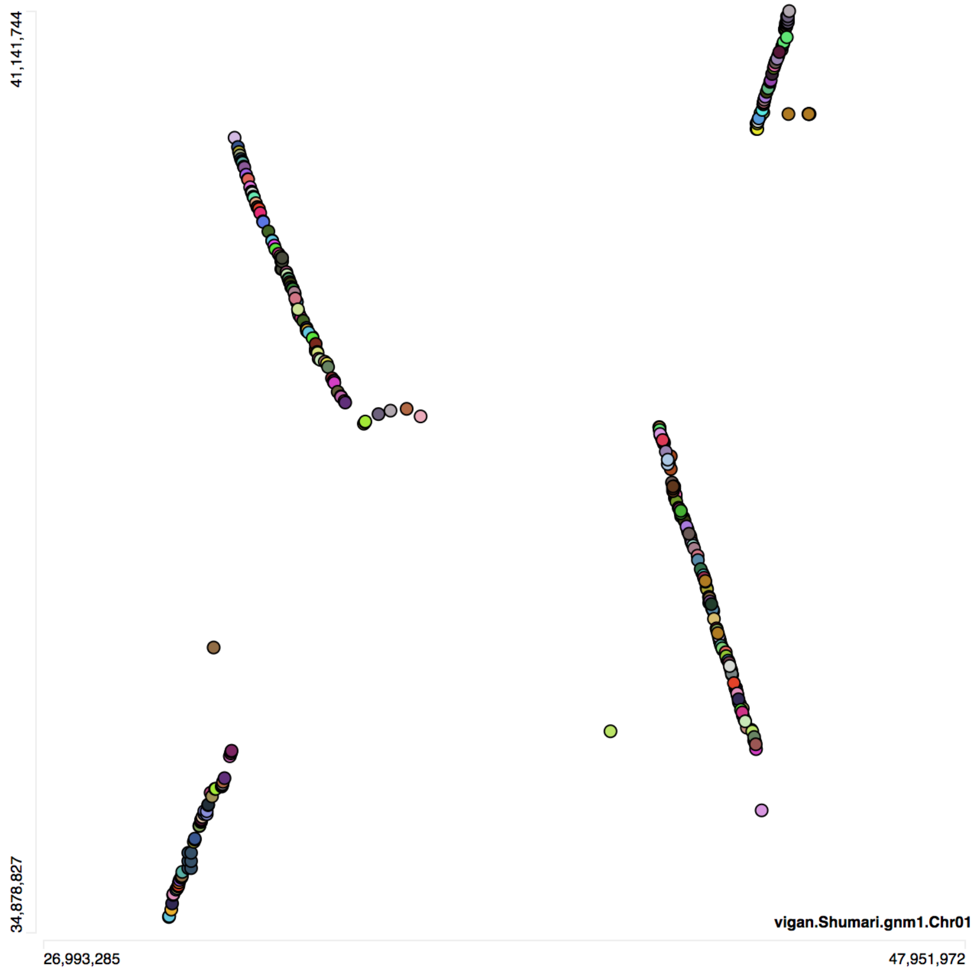

**Supplementary Figure 12.** Histogram obtained after flow cytometric analysis of prodium-iodide stained suspensions of cell nuclei isolated from *V. unguiculata* and *S. lycopersicum*. While peaks representing nuclei in G1 phase of cell cycle are clearly visible, peaks representing nuclei in G2 phase are small, indicating the presence of only a small fraction of cycling cells and/or cells arrested in G2 phase in leaf tissues. Average G1 peak positions were 100.3 and 150.1 for *V. unguiculata* and *S. lycopersicum*, respectively.

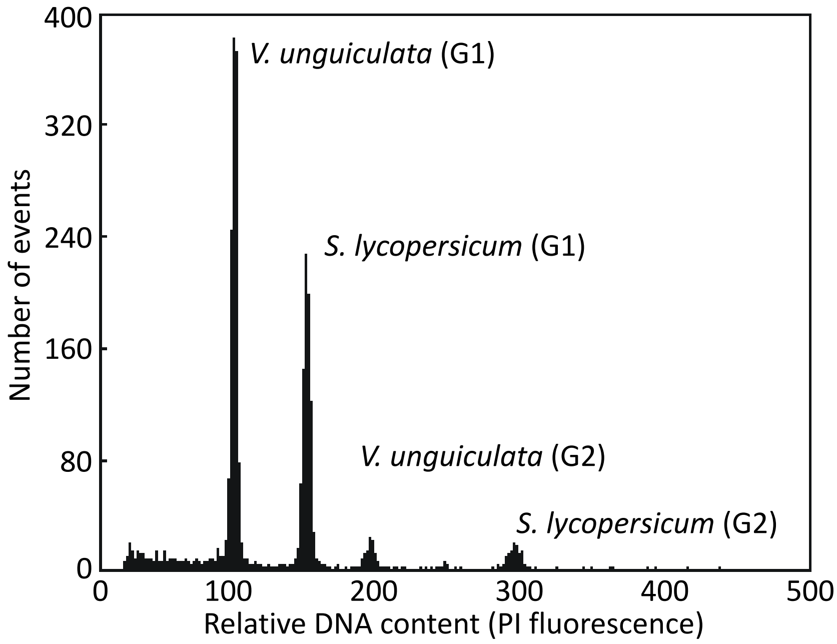

**Supplementary Table 1.** Statistics for *BspQI* optical map.

| | |
|---|---|
| # Molecules | 230,645 |
| Molecule N50 | 340.5 Kb |
| Molecule minimum length | 180 Kb |
| Molecule total length | 78.5 Gb |
| # BNG (optical map) contigs | 508 |
| BNG (optical map) total length | 622.21 Mb |
| BNG (optical map) N50 | 1.62 Mb |

**Supplementary Table 2.** Statistics for the *BssSI* optical map.

| | |
|---|---|
| # Molecules | 242,927 |
| Molecule N50 | 294.9 Kb |
| Molecule minimum length | 180 Kb |
| Molecule total length | 71.5 Gb |
| # BNG (optical map) contigs | 743 |
| BNG (optical map) total length | 577.76 Mb |
| BNG (optical map) N50 | 1.02 Mb |

**Supplementary Table 3.** Assembly statistics for the eight individual draft assemblies.

| | **CANU[a]** | **CANU[b]** | **ABruijn** | **FALCON** | **CANU[c]** | **CANU[d]** | **CANU[e]** | **CANU[f]** |
|---|---|---|---|---|---|---|---|---|
| Input reads (Gbp) | 56.6 | 56.6 | N/A | N/A | 56.6 | 56.6 | 56.6 | 56.6 |
| Input reads corrected (Gbp) | 17.5 | 30.6 | 30.6 | 30.6 | 15.9 | 19.5 | 18.8 | 31.6 |
| Input reads corrected (fold) | 28.2x | 49.4x | 49.4x | 49.4x | 25.7x | 31.7x | 30.6x | 51.6x |
| N50 (bp) | 5,307,785 | 4,754,622 | 2,157,242 | 3,138,839 | 3,751,474 | 3,175,625 | 2,798,135 | 5,641,635 |
| L50 | 27 | 31 | 67 | 45 | 41 | 47 | 48 | 28 |
| NG50 (bp) | 3,966,144 | 3,849,144 | 1,510,173 | 2,084,336 | 2,758,179 | 2,303,516 | 2,264,855 | 4,070,253 |
| LG50 | 39 | 43 | 107 | 65 | 59 | 66 | 69 | 39 |
| total (bp) | 505,857,042 | 517,175,664 | 478,230,679 | 511,933,729 | 504,711,938 | 516,558,510 | 515,964,327 | 510,383,709 |
| contigs | 878 | 946 | 498 | 1,789 | 1,053 | 1,093 | 1,119 | 944 |
| contigs ≥100kbp | 200 | 277 | 399 | 376 | 281 | 338 | 313 | 265 |
| contigs ≥ 1Mbp | 98 | 103 | 149 | 114 | 123 | 133 | 139 | 94 |
| contigs ≥ 10Mp | 10 | 10 | 1 | 1 | 2 | 4 | 3 | 10 |
| longest contig (bp) | 18,457,871 | 18,479,951 | 12,614,652 | 10,554,495 | 14,090,735 | 14,493,815 | 13,236,031 | 17,184,516 |
| mapped SNPs | 49,804 | 49,863 | 49,710 | 49,587 | 49,847 | 49,830 | 49,823 | 49,863 |

**N50:** length for which the set of contigs of that length or longer accounts for at least half of the assembly size
**NG50:** length for which the set of contigs of that length or longer accounts for at least half of the ~620Mb genome
**L50:** minimum number of contigs accounting for at least half of the assembly
**LG50:** minimum number of contigs accounting for at least half of the ~620Mb genome
**Assembly parameters:** assembly tools were run with default parameters, except for the following CANU[a] (corMhapSensitivity=low), CANU[b] (corMhapSensitivity=low, corOutCoverage=100), ABruijn (k=19, cov=49), FALCON (default), CANU[c] (corMhapSensitivity=high), CANU[d] (corMhapSensitivity=high, corMaxEvidenceErate=0.15, corOutCoverage=100), CANU[e] (corMhapSensitivity=normal, corMaxEvidenceErate=0.15, corOutCoverage=100), CANU[f] (corMhapSensitivity=high corOutCoverage=100)
**Mapped SNPs:** number of SNP design sequenced mapped to each assembly with an e-score of 1e-50 or better (see text)

**Supplementary Table 4.** Characteristics of the 10 genetic maps used for pseudomolecule construction. Cowpea chromosomes are numbered based on the new system (cross-reference shown in Supplementary Table 5).

| Genetic map | Characteristic | Vu01 | Vu02 | Vu03 | Vu04 | Vu05 | Vu06 | Vu07 | Vu08 | Vu09 | Vu10 | Vu11 | All |
|---|---|---|---|---|---|---|---|---|---|---|---|---|---|
| Tvu-14676 x IT84S-2246-4[5] | Markers | 1,207 | 953 | 2,765 | 1,066 | 1,563 | 1,326 | 1,447 | 1,618 | 1,250 | 774 | 691 | 14,660 |
| | Bins | 111 | 82 | 186 | 81 | 126 | 119 | 105 | 131 | 126 | 67 | 82 | 1,216 |
| | cM | 66.96 | 68.74 | 121.29 | 55.80 | 84.50 | 70.46 | 56.85 | 67.14 | 94.18 | 56.57 | 70.41 | 812.90 |
| Sanzi x Vita7[5] | Markers | 1,256 | 1,376 | 2,912 | 1,393 | 1,156 | 973 | 1,920 | 1,050 | 1,343 | 851 | 1,389 | 15,619 |
| | Bins | 115 | 77 | 215 | 103 | 92 | 85 | 157 | 112 | 116 | 83 | 110 | 1,265 |
| | cM | 77.59 | 50.97 | 156.40 | 84.95 | 82.07 | 70.13 | 101.01 | 85.03 | 100.46 | 66.37 | 80.51 | 955.51 |
| ZN016 x Zhijiang282[5] | Markers | 857 | 580 | 1,551 | 858 | 800 | 690 | 426 | 562 | 791 | 456 | 393 | 7,964 |
| | Bins | 55 | 57 | 123 | 49 | 70 | 60 | 49 | 56 | 83 | 43 | 52 | 697 |
| | cM | 71.49 | 65.27 | 124.01 | 45.09 | 91.70 | 62.88 | 54.48 | 75.47 | 94.07 | 56.64 | 62.28 | 803.38 |
| CB46 x IT93K-503-1[5] | Markers | 1,374 | 1,138 | 2,745 | 855 | 1,342 | 1,151 | 2,050 | 1,336 | 1,759 | 1,601 | 1,227 | 16,578 |
| | Bins | 88 | 92 | 179 | 66 | 109 | 74 | 109 | 91 | 116 | 83 | 76 | 1,083 |
| | cM | 73.09 | 67.87 | 132.85 | 56.00 | 94.54 | 57.64 | 84.46 | 67.26 | 85.50 | 62.87 | 59.77 | 841.84 |
| CB27 x IT82E-18[5] | Markers | 737 | 1,534 | 2,640 | 1,843 | 1,520 | 1,539 | 1,295 | 1,255 | 1,195 | 1,778 | 1,230 | 16,566 |
| | Bins | 37 | 68 | 159 | 93 | 100 | 62 | 100 | 105 | 109 | 69 | 75 | 977 |
| | cM | 59.07 | 46.54 | 132.64 | 82.06 | 95.31 | 60.01 | 78.47 | 90.07 | 86.69 | 59.76 | 68.30 | 858.91 |
| CB27 x IT97K-566-6 | Markers | 1,577 | 1,205 | 3,174 | 1,230 | 796 | 1,576 | 1,212 | 803 | 1,721 | 1,735 | 1,255 | 16,284 |
| | Bins | 144 | 59 | 144 | 38 | 78 | 68 | 97 | 52 | 117 | 52 | 72 | 921 |
| | cM | 86.47 | 47.87 | 126.36 | 56.92 | 83.12 | 57.04 | 88.59 | 61.01 | 92.39 | 51.88 | 76.70 | 828 |
| 524B x IT84S-2049[27] | Markers | 738 | 981 | 2,483 | 1,234 | 942 | 1,425 | 1,257 | 1,311 | 1,626 | 961 | 1,244 | 14,202 |
| | Bins | 43 | 72 | 175 | 84 | 66 | 64 | 105 | 97 | 114 | 74 | 57 | 951 |
| | cM | 69.72 | 60.44 | 148.52 | 86.08 | 71.80 | 63.93 | 105.19 | 76.80 | 110.95 | 64.11 | 51.99 | 910 |
| CB46-Null x FN-2-9-04 | Markers | 1,168 | 1,454 | 3,354 | 1,245 | 1,132 | 1,342 | 1,798 | 1,407 | 1,678 | 961 | 1,669 | 17,208 |
| | Bins | 126 | 106 | 250 | 102 | 82 | 108 | 147 | 121 | 151 | 91 | 108 | 1,392 |
| | cM | 80.94 | 74.72 | 150.83 | 92.78 | 60.02 | 65.30 | 110.66 | 81.61 | 107.51 | 76.27 | 85.25 | 986 |
| CB27 x UCR779 | Markers | 1,565 | 1,475 | 3,684 | 1,481 | 1,103 | 1,556 | 1,792 | 1,635 | 1,519 | 772 | 1,632 | 18,214 |
| | Bins | 52 | 45 | 111 | 47 | 34 | 91 | 66 | 48 | 63 | 39 | 48 | 644 |
| | cM | 60.18 | 61.36 | 125.55 | 58.17 | 76.37 | 62.91 | 84.60 | 54.58 | 75.75 | 70.22 | 56.04 | 786 |
| IT99K-573-1-1 x TVNu-1158[9] | Markers | 1,742 | 1,368 | 2,780 | 117 | 1,740 | 1,676 | 1,968 | 1,607 | 1,774 | 1,416 | 1,551 | 17,739 |
| | Bins | 132 | 140 | 280 | 24 | 205 | 150 | 228 | 159 | 189 | 144 | 174 | 1,825 |
| | cM | 65.31 | 98.62 | 157.61 | 15.25 | 114.62 | 78.29 | 134.15 | 81.59 | 100.37 | 77.37 | 102.85 | 1,026 |

**Supplementary Table 5.** Cross-reference between old and new chromosome numbers for cowpea (Vu). The total length of the syntenic matches (exact match >100 bp, alignment length > 1 kb) with the top two *P. vulgaris* (Pv) chromosomes is shown. Chromosomes that were inverted to meet the “short arm on top” convention are indicated in parenthesis. (*) Optimal solution.

| Old Vu Chr. | Pv Chr (kb) | Pv Chr (kb) | New Vu Chr. |
|---|---|---|---|
| 1 | 8 (671.2) | **5** (485.1) | **5*** |
| 2 | **7** (1390.0) | | **7** |
| 3 | **3** (1493.9) | 2 (899.8) | **3** |
| 4 | **1** (932.0) | 5 (245.0) | **1** |
| 5 | **8** (573.3) | 1 (309.0) | **8*** |
| 6 | **6** (996.8) | | **6** (inverted) |
| 7 | **2** (736.9) | 3 (163.8) | **2** |
| 8 | **9** (1439.5) | | **9** |
| 9 | **11** (751.8) | | **11** (inverted) |
| 10 | **10** (593.9) | | **10** (inverted) |
| 11 | **4** (564.1) | | **4** |

**Supplementary Table 6.** Annotated repeat abundances in cowpea. The major represented classes, super-families, and subgroups of transposable elements as determined by automated annotation and classified according to the scheme of Wicker et al.[29], as well as other major repeat types are presented.

| | % of genome | % of TE (bp) | Number | Number (%) | Sum (Mbp) | Average length (bp) |
|---|---|---|---|---|---|---|
| **All repeats** | **49.53** | | | | | |
| **Mobile Element** | **39.23** | **100.00** | | **100.00** | | |
| **Class I: Retroelement (RXX)** | **33.17** | **84.55** | **241542** | **82.32** | **179.035** | |
| LTR Retrotransposon (RLX) | 32.76 | 83.49 | 230558 | 80.92 | 170.146 | |
| *Gypsy* (RLG) | 18.32 | 46.70 | 97313 | 34.17 | 95.162 | 978 |
| *Copia* (RLC) | 11.80 | 30.09 | 107705 | 37.80 | 61.317 | 569 |
| TRIMs | 0.002 | 0.01 | 57 | 0.02 | 0.012 | 206 |
| unclassified LTR (RLX) | 2.48 | 6.31 | 24637 | 8.65 | 12.860 | 522 |
| non-LTR Retrotransposon (RXX) | 0.41 | 1.06 | 3971 | 1.39 | 2.153 | |
| LINE (RIX) | 0.36 | 0.92 | 3571 | 1.25 | 1.883 | 527 |
| SINE (RSX) | 0.05 | 0.13 | 400 | 0.14 | 0.269 | 673 |
| **Class II: DNA Transposon (DXX)** | **6.06** | **15.45** | **50376** | **17.68** | **31.491** | |
| DNA Transposon Superfamily (DTX) | 4.70 | 11.97 | 42902 | 15.06 | 24.402 | |
| CACTA (DTC) | 2.22 | 5.66 | 19538 | 6.86 | 11.538 | 591 |
| hAT (DTA) | 1.38 | 3.51 | 15684 | 5.50 | 7.159 | 456 |
| MuDR (DTM) | 0.94 | 2.40 | 6672 | 2.34 | 4.884 | 732 |
| MITE (DXX) | 0.05 | 0.11 | 218 | 0.08 | 0.233 | 1067 |
| Helitron (DHH) | 1.30 | 3.31 | 7013 | 2.46 | 6.736 | 961 |
| unclassified DNA transposon (DXX) | 0.02 | 0.06 | 239 | 0.08 | 0.119 | 498 |
| **Class I/Class II ratio** | **7.23** | | **5.57** | | | |
| ***Gypsy/Copia* ratio** | **1.55** | | **0.90** | | | |
| **Other repeats** | | | | | | |
| Simple Sequence Repeats (SSRs) | 4.03 | | 188510 | | 20. 936 | 111 |
| Satellite repeats | 0.06 | | 93 | | 0.330 | 3551 |
| Ribosomal DNA (rDNA) | 0.49 | | 505 | | 2. 555 | |
| Unknown low-complexity sequence | 5.68 | | 76875 | | 29.515 | |

**Supplementary Table 7.** Start and end positions of centromeric regions based on BLAST of the 455-bp tandem repeat identified by Iwata-Otsubo et al.[26].

| Chromosome | Start (bp) | End (bp) | Range (bp) |
|---|---|---|---|
| Vu01 | 14,698,036 | 16,525,496 | 1,827,460 |
| Vu02 | 10,238,236 | 14,020,258 | 3,782,022 |
| Vu03 | 30,476,981 | 31,470,261 | 993,280 |
| Vu04 | 19,069,641 | 21,130,843 | 2,061,202 |
| Vu05 | 25,704,431 | 33,885,354 | 8,180,923 |
| Vu06 | 9,156,830 | 9,235,637 | 78,807 |
| Vu07 | 16,587,031 | 16,604,960 | 17,929 |
| Vu08 | 14,914,119 | 15,164,402 | 250,283 |
| Vu09 | 20,802,610 | 22,685,597 | 1,882,987 |
| Vu10 | 18,917,563 | 19,028,450 | 110,887 |
| Vu11 | 17,283,961 | 18,283,861 | 999,900 |

**Supplementary Table 10.** Number and location of SNPs relative to annotated cowpea genes.

| | **1M list** | **iSelect** |
|---|---|---|
| # SNPs | 957,710 | 51,128 |
| # SNPs in genes (%) | 336,285 (35%) | 31,708 (62%) |
| # SNPs in exons (%) | 138,892 (15%) | 16,898 (33%) |
| # SNPs in or within 1 kb from gene (%) | 460,709 (48%) | 38,286 (75%) |
| # SNPs in or within 2 kb from gene (%) | 540,773 (56%) | 39,856 (78%) |
| # SNPs in or within 10 kb from gene (%) | 792,318 (83%) | 45,648 (89%) |
| # unique genes with SNPs (%) | 23,266 (78%) | 17,444 (59%) |
| # unique genes containing or near SNPs (< 1 kb) (%) | 25,433 (85%) | 19,319 (65%) |
| # unique genes containing or near SNPs (< 2 kb) (%) | 26,130 (88%) | 19,818 (67%) |
| # unique genes containing or near SNPs (< 10 kb) (%) | 27,021 (91%) | 21,205 (71%) |

**Supplementary Table 15.** Comparative repeat abundance in *Vigna* species. Repeats in each genome assembly were annotated by common methods and categorized into the major groups as in Supplementary Table 6.

| | ***V. unguiculata* (% genome)** | ***V. angularis* (% genome)** | ***V. radiata* (% genome)** | **Vu vs. Vr (Mbp)** | **Vu vs. Va (Mbp)** | **Va vs. Vr (Mbp)** |
|---|---|---|---|---|---|---|
| **Genome assembly size (Mbp)** | 519.44 | 467.30 | 463.64 | 55.798 | 52.135 | 3.663 |
| **Mobile Element** | **39.23** | **34.21** | **32.69** | **52.236** | **43.907** | **8.330** |
| **Class I: Retroelement (RXX)** | **33.17** | **31.93** | **30.13** | **32.599** | **23.113** | **9.486** |
| LTR Retrotransposon (RLX) | 32.76 | 31.68 | 29.83 | 31.836 | 22.103 | 9.733 |
| *Gypsy* (RLG) | 18.32 | 15.12 | 14.02 | 30.155 | 24.525 | 5.631 |
| *Copia* (RLC) | 11.80 | 11.68 | 11.49 | 8.061 | 6.723 | 1.338 |
| unclassified LTR (RLX) | 2.48 | 4.85 | 4.22 | -6.692 | -9.801 | 3.108 |
| non-LTR Retrotransposon (RXX) | 0.41 | 0.12 | 0.12 | 1.577 | 1.584 | -0.008 |
| LINE (RIX) | 0.36 | 0.11 | 0.12 | 1.343 | 1.362 | -0.019 |
| SINE (RSX) | 0.05 | 0.01 | 0.01 | 0.234 | 0.223 | 0.011 |
| **Class II: DNA Transposon (DXX)** | **6.06** | **2.29** | **2.16** | **21.457** | **20.794** | **0.664** |
| DNA Transposon Superfamily (DTX) | 4.70 | 2.19 | 2.10 | 14.675 | 14.161 | 0.514 |
| CACTA (DTC) | 2.22 | 1.29 | 0.98 | 6.972 | 5.497 | 1.475 |
| hAT (DTA) | 1.38 | 0.33 | 0.43 | 5.170 | 5.622 | -0.452 |
| MuDR (DTM) | 0.94 | 0.50 | 0.60 | 2.100 | 2.555 | -0.455 |
| Helitron (DHH) | 1.30 | 0.12 | 0.17 | 5.927 | 6.162 | -0.235 |
| **Class I/Class II ratio** | **7.2** | **13.9** | **13.9** | | | |
| ***Gypsy*/*Copia* ratio** | **1.55** | **1.22** | **1.29** | | | |
| **Other repeats** | | | | | | |
| Simple Sequence Repeats (SSRs) | 4.03 | 2.25 | 1.50 | 0.835 | -2.722 | 3.557 |

**Supplementary Table 17.** Information of the PCR primer sequences designed to amplify the two breakpoints of the inversion using both the orientation of the reference and the opposite orientation.

| Primer | Sequence | Tm (°C) | Amplicon size (bp) | Breakpoint target | Orientation |
|---|---|---|---|---|---|
| BP1_Ref_Forward | CCTTGTCCCCCATTTTTCTT | 60.16 | | 1 | Reference |
| BP1_Ref_Reverse | TGATGTGAAATTGTGATCCATGT | 60.11 | 822 | 1 | Reference |
| BP1_Opp_Forward | CCTTGTCCCCCATTTTTCTT | 60.16 | | 1 | Opposite |
| BP1_Opp_Reverse | TTGAGCACCAAAGTGTCGAA | 60.43 | 796 | 1 | Opposite |
| BP2_Ref_Forward | AAAATGACCGGAATCATGAAC | 58.75 | | 2 | Reference |
| BP2_Ref_Reverse | TTACCATTCGCAACGAAAAA | 59.19 | 389 | 2 | Reference |
| BP2_Opp_Forward | TGATGTGAAATTGTGATCCATGT | 60.11 | | 2 | Opposite |
| BP2_Opp_Reverse | TTACCATTCGCAACGAAAAA | 59.19 | 822 | 2 | Opposite |

**Supplementary Table 18.** Data sources and references for genome assemblies and annotations used in the gene family analysis.

| Genus species | Abbreviation | Genotype | Assembly | Annotation | Citation | Source | Original filename |
|---|---|---|---|---|---|---|---|
| *Vigna unguiculata* | vigun | IT97K-499-35 | 1 | 1 | This study | Phytozome | Vunguiculata_469_v1.1.protein_primaryTranscriptOnly.fa |
| *Vigna angularis* | vigan | Shumari | 1 | 1 | 74 | VigGS | Vangularis_v1.a1.protein.fasta |
| *Vigna radiata* | vigra | VC1973A | 6 | 1 | 39 | LegumeInfo | vigra.VC1973A.gnm6.ann1.M1Qs.protein.faa |
| *Phaseolus vulgaris* | phavu | G19833 | 2 | 1 | 23 | Phytozome | Pvulgaris_442_v2.1.protein.fa |
| *Cajanus cajan* | cajca | ICPL87119 | 1 | 1 | 24 | LegumeInfo | cajca.ICPL87119.gnm1.ann1.Y27M.protein_main.faa |
| *Glycine max* | glyma | Williams 82 | 2 | 1 | 30 | Phytozome | Gmax_275_Wm82.a2.v1.protein_primaryTranscriptOnly.fa |
| *Medicago truncatula* | medtr | A17_HM341 | 4 | 2 | 83 | LegumeInfo | medtr.A17_HM341.gnm4.ann2.G3ZY.pep.faa |
| *Cicer arietinum* | cicar | Frontier | 1 | 1 | 84 | LegumeInfo | cicar.CDCFrontier.gnm1.ann1.nRhs.gene.pep.faa |
| *Trifolium pratense* | tripr | MilvusB | 2 | 1 | 85 | LegumeInfo | tripr.MilvusB.gnm2.ann1.DFgp.protein_primaryTranscript.faa |
| *Lotus japonicus* | lotja | MG20 | 3 | 1 | 86 | LegumeInfo | lotja.MG20.gnm3.QPGB.protein.faa |
| *Lupinus angustifolius* | lupan | Tanjil | 1 | 1 | 87 | LegumeInfo | lupan.Tanjil.gnm1.ann1.nnV9.protein_all.faa |
| *Arachis duranensis* | aradu | V14167 | 1 | 1 | 88 | LegumeInfo | aradu.V14167.gnm1.ann1.cxSM.protein.faa |
| *Arachis ipaensis* | araip | K30076 | 1 | 1 | 88 | LegumeInfo | araip.K30076.gnm1.ann1.J37m.protein.faa |
| *Arachis hypogaea* | arahy | Tifrunner | 1 | 1 | Peanutbase | LegumeInfo | arahy.Tifrunner.gnm1.ann1.CCJH.protein_primaryTranscript.faa |
| *Arabidopsis thaliana* | arath | Col-0 | TAIR10 | 1 | TAIR | Phytozome | Athaliana_167_TAIR10.protein_primaryTranscriptOnly.fa |
| *Cucumis sativus* | cucsa | unknown | 1 | 1 | Phytozome | Phytozome | Csativus_122_v1.0.protein_primaryTranscriptOnly.fa |
| *Prunus persica* | prupe | Lovell | 2 | 2.1 | 89 | Phytozome | Ppersica_298_v2.1.protein_primaryTranscriptOnly.fa |
| *Solanum lycopersicum* | solly | Heinz_1706 | 2.5 | ITAG2.4 | 90 | Phytozome | Slycopersicum_390_ITAG2.4.protein_primaryTranscriptOnly.fa |
| *Vitis vinifera* | vitvi | PN40024 | Genoscope.12X | Genoscope.12X | 91 | Phytozome | Vvinifera_145_Genoscope.12X.protein_primaryTranscriptOnly.fa |
